# Multiple spillovers and onward transmission of SARS-CoV-2 in free-living and captive white-tailed deer

**DOI:** 10.1101/2021.10.31.466677

**Authors:** Suresh V. Kuchipudi, Meera Surendran-Nair, Rachel M. Ruden, Michelle Yon, Ruth H. Nissly, Rahul K. Nelli, Lingling Li, Bhushan M. Jayarao, Kurt J. Vandegrift, Costas D. Maranas, Nicole Levine, Katriina Willgert, Andrew J. K. Conlan, Randall J. Olsen, James J. Davis, James M. Musser, Peter J. Hudson, Vivek Kapur

## Abstract

Many animal species are susceptible to SARS-CoV-2 and could potentially act as reservoirs, yet transmission of the virus in non-human free-living animals has not been documented. White-tailed deer (*Odocoileus virginianus*), the predominant cervid in North America, are susceptible to SARS-CoV-2 infection, and experimentally infected fawns can transmit the virus. To test the hypothesis that SARS-CoV-2 may be circulating in deer, we tested 283 retropharyngeal lymph node (RPLN) samples collected from 151 free-living and 132 captive deer in Iowa from April 2020 through December of 2020 for the presence of SARS-CoV-2 RNA. Ninety-four of the 283 deer (33.2%; 95% CI: 28, 38.9) samples were positive for SARS-CoV-2 RNA as assessed by RT-PCR. Notably, between November 23, 2020 and January 10, 2021, 80 of 97 (82.5%; 95% CI 73.7, 88.8) RPLN samples had detectable SARS-CoV-2 RNA by RT-PCR. Whole genome sequencing of the 94 positive RPLN samples identified 12 SARS-CoV-2 lineages, with B.1.2 (n = 51; 54.5%), and B.1.311 (*n* = 19; 20%) accounting for ~75% of all samples. The geographic distribution and nesting of clusters of deer and human lineages strongly suggest multiple zooanthroponotic spillover events and deer-to-deer transmission. The discovery of sylvatic and enzootic SARS-CoV-2 transmission in deer has important implications for the ecology and long-term persistence, as well as the potential for spillover to other animals and spillback into humans. These findings highlight an urgent need for a robust and proactive “One Health” approach to obtaining a better understanding of the ecology and evolution of SARS-CoV-2.

**One-Sentence Summary:** SARS-CoV-2 was detected in one-third of sampled white-tailed deer in Iowa between September 2020 and January of 2021 that likely resulted from multiple human-to-deer spillover and deer-to-deer transmission events.

## Introduction

Severe acute respiratory syndrome coronavirus 2 (SARS-CoV-2), the cause of coronavirus disease 2019 (COVID-19) in humans, is a novel coronavirus in the genus *Betacoronavirus* (subgenus *Sarbecovirus*) (1). SARS-CoV-2 was first identified in Wuhan, China, toward the end of 2019 (2), and has caused a pandemic with 247 million COVID-19 cases and over 5 million deaths globally as of November 1^st^, 2021 (3). The virus continues to evolve, with a growing concern for the emergence of new variants. SARS-CoV-2 uses the host angiotensin-converting enzyme 2 (ACE-2) receptor to enter cells (4). ACE-2 receptors are well conserved across vertebrate species including humans (5), and computational analyses predict high binding affinities of SARS-CoV-2 to the ACE-2 receptor in multiple animal species, indicating potential susceptibility to infection (5). Included amongst these are three species of cervids: the Père David’s deer (*Elaphurus davidianus*), reindeer (*Rangifer tarandus*), and white-tailed deer (*Odocoileus virginianus*) (6).

The widespread and global dissemination of SARS-CoV-2 among humans provides opportunities for spillovers into non-human hosts (7). Indeed, SARS-CoV-2 infections have been documented in dogs, cats, zoo animals (e.g. tigers and lions) and farmed mink (8, 9). In principle, SARS-CoV-2 infection of a non-human animal host might result in the establishment of a reservoir that can further drive the emergence of novel variants with potential for spillback to humans. This type of transmission cycle has been described among workers on mink farms (10). However, widespread SARS-CoV-2 transmission in a free-living animal species has not yet been documented.

Our study was prompted by a recent report that 40% of free-living white-tailed deer in the USA had antibodies against SARS-CoV-2 (11). More recent studies have provided evidence of SARS-CoV-2 transmission among experimentally infected deer under controlled conditions (6). To test the hypothesis that infection and subsequent transmission of SARS-CoV-2 of deer occurs in nature, we assayed 283 retropharyngeal lymph node (RPLN) samples collected from free-living and captive deer in Iowa from April 2020 through January 2021. We discovered that one-third of the deer sampled over the course of the study had SARS-CoV-2 nucleic acid in their RPLN. We then sequenced the SARS-CoV-2 genomes present in all positive samples and found that the genomes represented multiple lineages that corresponded to viral genotypes circulating contemporaneously in humans. In the aggregate, our results are consistent with a model of multiple independent human-to-deer transmission events and deer-to-deer transmission. Our findings raise the possibility of reverse zoonoses, especially in exurban areas with high deer density. The study also highlights the potential risks and considerable knowledge gaps associated with the continued evolution of SARS-CoV-2 in animal hosts, and calls for the implementation of enhanced surveillance programs to identify potential reservoir species at the animal-human interface.

## Results and Discussion

### SARS-CoV-2 RNA is present in a substantial fraction of free-living and captive deer across Iowa, with strong evidence of temporal clustering

A total of 283 RPLN samples recovered from wild and captive deer in Iowa between April 8, 2020 and January 6, 2021 were analyzed by RT-PCR for the presence of SARS-CoV-2 RNA signatures using the OPTI Medical SARS-CoV-2 RT-PCR assay targeting the N1 and N2 genes in a CLIA-approved laboratory ((12); Supplementary Table 1). The sampling period closely follows the trajectory of the pandemic in Iowa (with the first human case reported on March 8, 2020) through the peak during the second week of November 2020 ((3); Figure 1).

**Figure 1:**
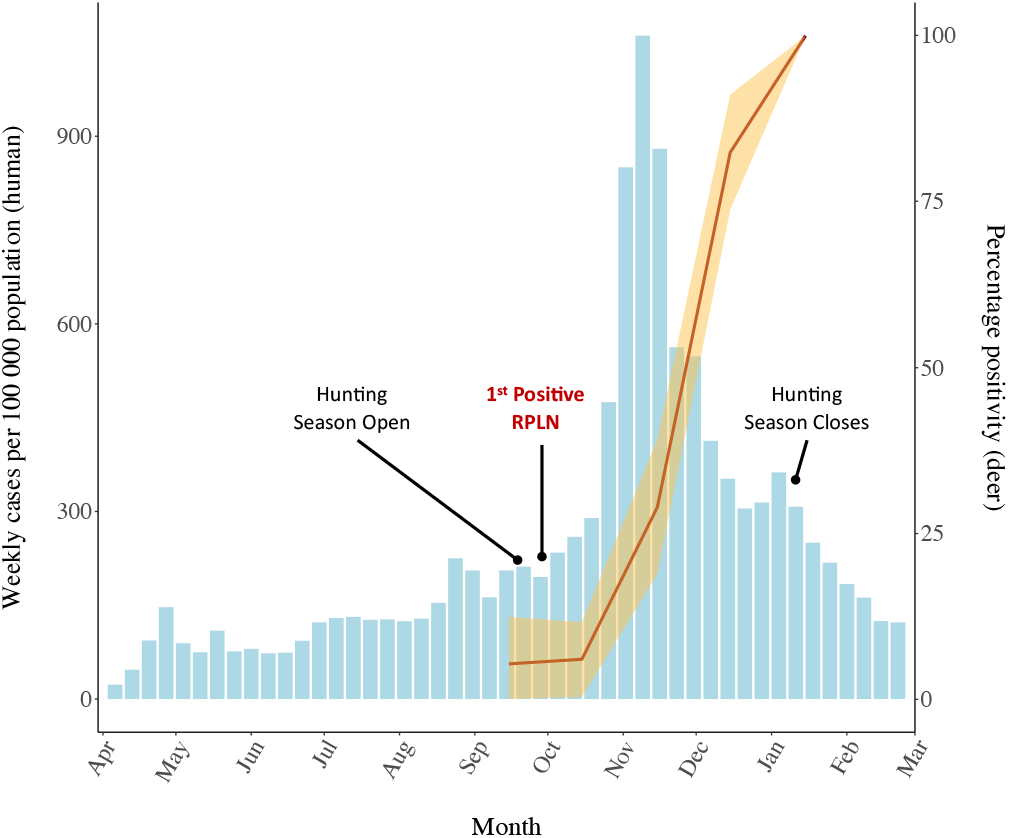
Epidemic curve showing SARS-CoV-2 weekly cases (per 100,000) in humans and the monthly change in SARS-CoV-2 positivity in white-tailed deer in Iowa. The histogram represents the progression in weekly reported cases (per 100,000 individuals; left axis). The percentage (and 95%) CI of white-tailed deer found positive for the presence of SARS-CoV-2 RNA (red line; right axis) appears to closely follow the trajectory of the human pandemic. The first identified positive sample on September 28, 2020 is marked, as is the start of the regular hunting season on September 19, 2020 and its end on January 10, 2021.

Sources and characteristics associated with the 283 RPLN samples are summarized in Table 1. Overall, 94 of the 283 (33.2%; 95% CI: 28, 38.9) RPLN samples were found to be positive for SARS-CoV-2 RNA. Since a majority (*n* = 261, 91.9%) of the 283 RPLN samples in our study were harvested from deer from September through December of 2020, a period that coincided with the regular deer hunting season in Iowa that started on September 19, 2020, and ended January 10, 2021, we explored temporal trends in the recovery of SARS-CoV-2 RNA from deer in the sample set. The results show that all 17 RPLNs from deer collected during April through August 2020 were negative for the presence of SARS-CoV-2 RNA (Table 1). The first identification of SARS-CoV-2 in deer was on September 28, 2020, in an animal harvested on a game preserve in South Eastern Iowa (Figure 2; Supplementary Table 1). This was closely followed by a second positive sample identified in a free-living deer killed in a road accident on September 30, 2020, approximately 300 miles away in Woodbury county on the State’s Western border (Supplementary Table 1). In total, two of the 39 samples collected in September were positive for SARS-CoV-2 RNA (5.1%: 95% CI: 1.4, 16.9). Similarly, in October 2020, four of 70 (5.7%; 95% CI 2.2, 13.8) RPLNs recovered from deer, one each from Des Moines and Pottawattamie counties, and two from Woodbury, were found to be positive for SARS-CoV-2 RNA. Coinciding with the major peak of infection in humans in Iowa, the positivity rate in deer rapidly increased with 22 of 77 RPLN samples (27.8%; 95% CI 19.2, 38.6) harboring SARS-CoV-2 RNA in November 2020, and 61 of 75 samples (81.3%; 95% CI: 71.1, 88.5) positive in December of 2020 (Figure 1). During the second week of January 2021, at the end of the regular hunting season, all five deer RPLN samples were positive for SARS-CoV-2 RNA (95% CI: 56.6-100). Notably, during the seven-weeks starting November 23, 2020, through the end of hunting season on January 10, 2021, 80 of 97 (82.5%; 95% CI 73.7, 88.8) deer RPLN samples from across the State were positive for SARS-CoV-2 RNA. Importantly, the results show a high viral copy number in deer RPLN samples (Median 106,000 viral copies per ml; ranging from 268 to 5.4 × 10^8^ copies per ml) (Supplementary Figure 1), suggesting that many of the deer likely had a very high viral load.

**Table 1:**
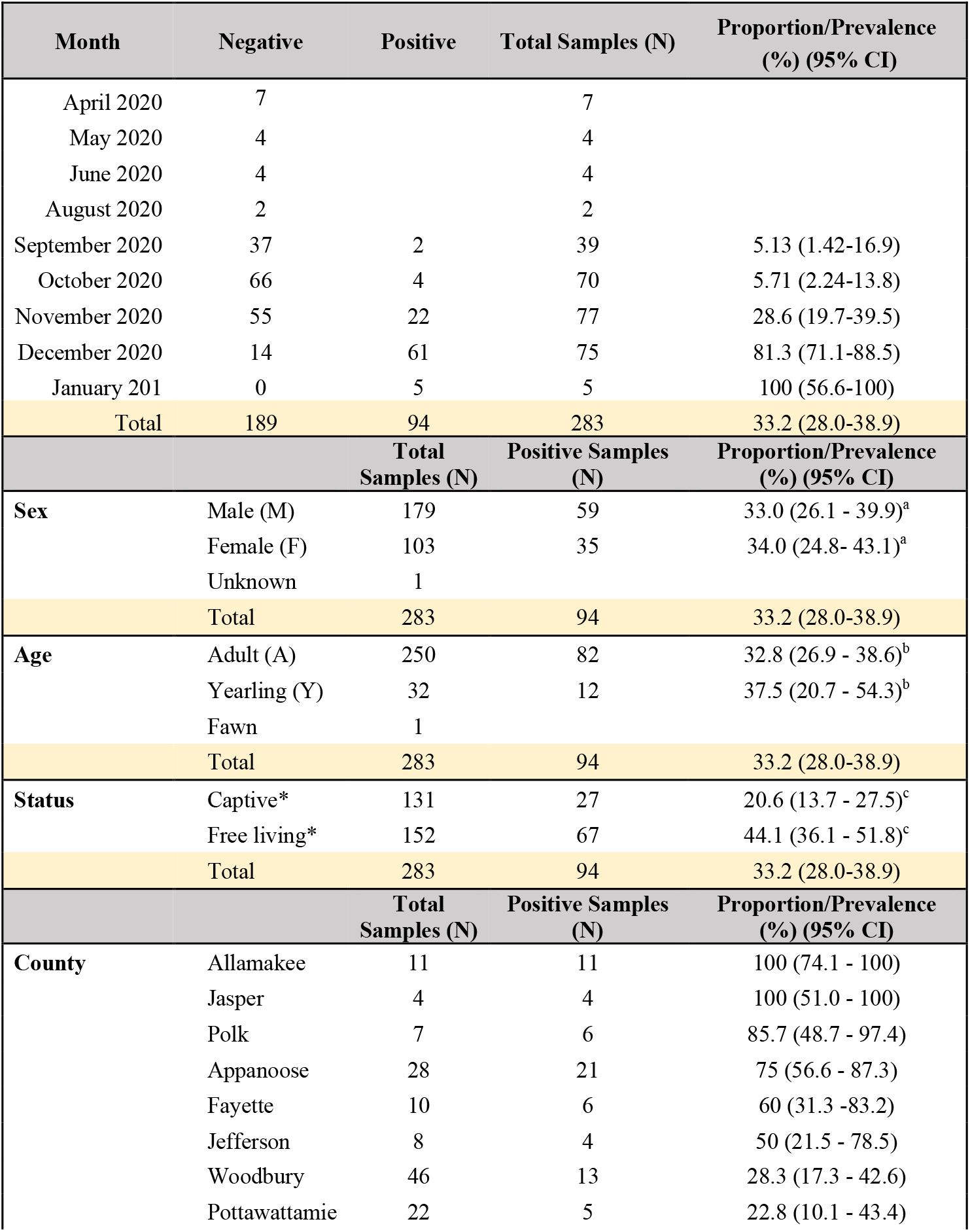
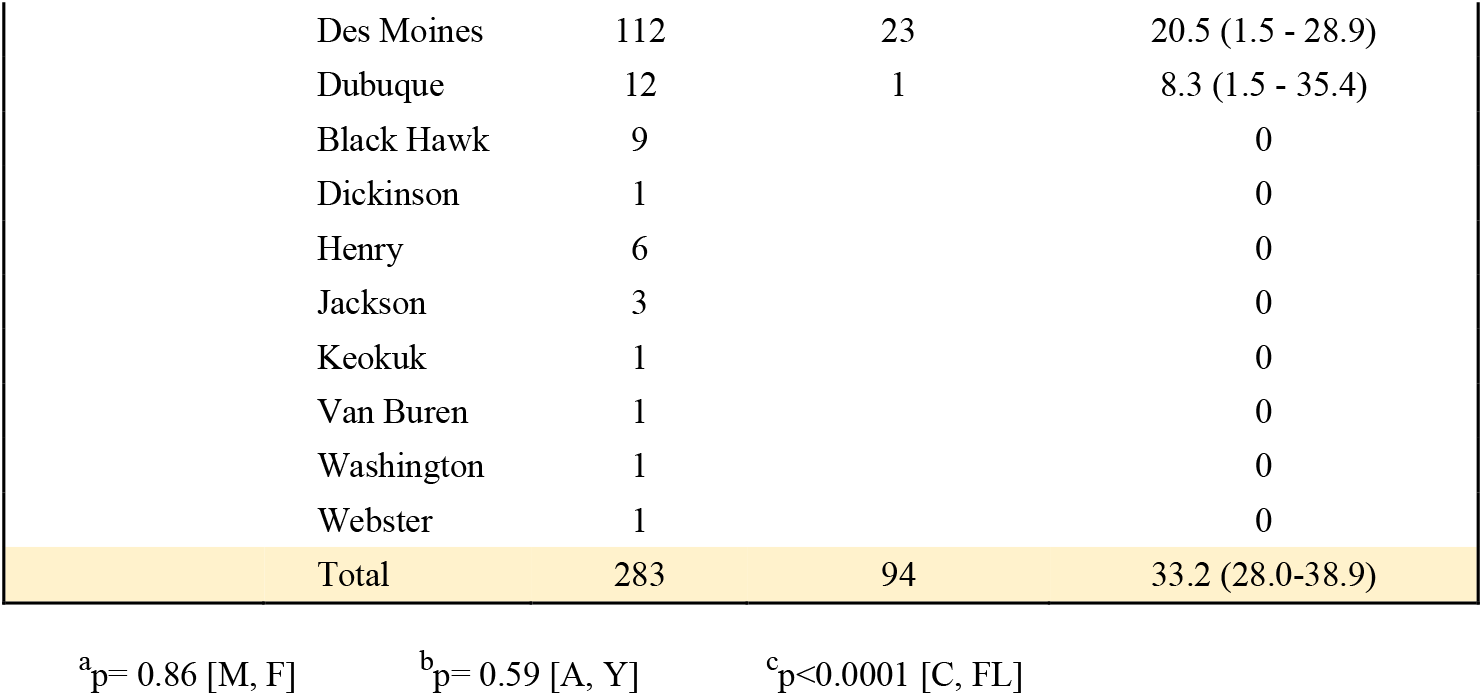
Characteristics and demographics of white-tailed deer screened for presence of SAR-CoV-2 in Iowa.

**Figure 2:**
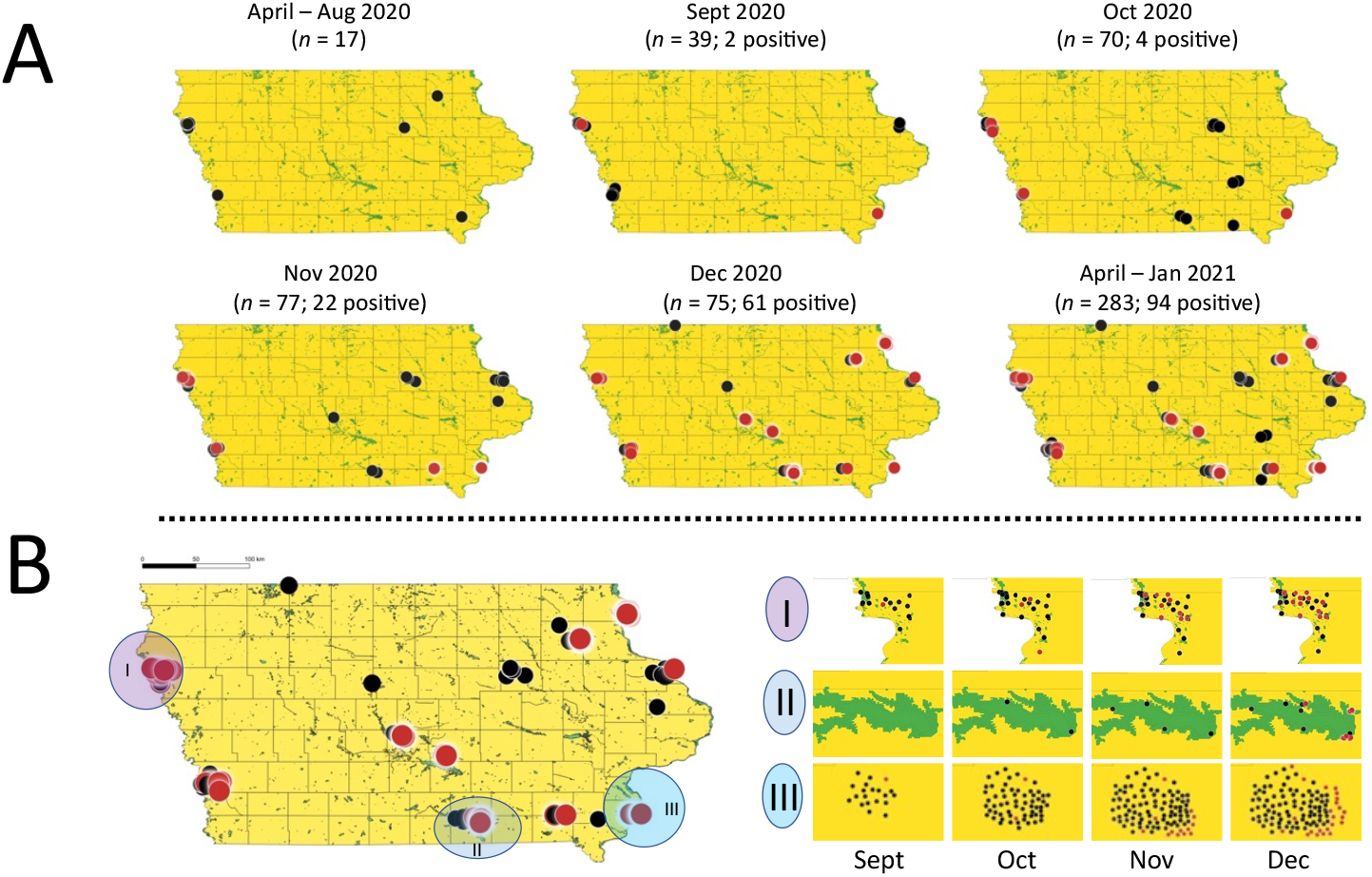
Temporal and spatial distribution of SARSCoV-2 positive samples from white-tailed deer in IA. The 94 SARS-CoV-2 positive deer RPLN samples from geographically dispersed sites were mapped and show strong temporal clustering in frequency of detection of positive samples starting with the first positive case identified in September. **A.** Monthly snapshots showing number and location of SARS-CoV-2 positive cases identified in deer in Iowa. **B.** Progression in number of positive cases in deer from September through December 2020 in three exemplar regions with different sampling intensities and sizes of sampling area. Each filled black circle represents a negative test result, and each filled red circle represents a positive test result for presence of SARS-CoV-2 RNA in deer RPLN.

### Ecological associations and risk factors for recovery of SARS-CoV-2 RNA from free-living and captive deer in Iowa

An exploratory analysis of potential risk factors for the identification of SARS-CoV-2 RNA in deer was carried out. We found no statistically significant differences (95% level) in the proportion of positive tests by age or sex within this sample (Table 1, Supplementary Figure 2). However, since this study was not designed or powered to probe this question with rigor, further investigations may be warranted.

The deer included in the study were either “free-living” on public lands or in peri-urban environments (*n* = 151) or were resident in nature or hunting preserves in “captive” settings (*n* = 132) (Table 1). Notably, the results suggest that the proportion of free-living deer positive for SARS-CoV-2 (44.4%; 95% CI: 36.4, 52.3) was significantly higher (z = 4.3; p < 0.0001) than in the deer living within preserves / captive settings (20.4%; 95% CI 13.7, 27.4). However, these results may be confounded in that four times as many RPLN were harvested from free-living deer (*n* = 52) as were from captive deer (*n* = 13) during December 2020, when the virus positivity rate was at its peak in Iowa deer herd. Hence, further studies are needed to assess the true significance or reasons for the observed differences in prevalence between free-living deer and that resident in managed environments. For instance, to better assess the risk of spillover and transmission, it is important to understand whether deer in managed settings are less stressed, have different nutritional status, or otherwise exhibit behaviors that influence opportunities for spillover or within-herd transmission of SARS-CoV-2 compared with those that are free-living.

We next explored regional differences in observed SARS-CoV-2 positivity among deer at the county level across the State. Figure 2A shows the widespread distribution of positive samples recovered from deer throughout Iowa and illustrates a strong temporal trend in SARS-CoV-2 positivity as the year progressed. The study identified 10 counties with at least one positive sample (Table 1). The largest number of RPLN samples represented in the collection were from a single game preserve (Preserve 2; Fig 2B) in Southeastern Iowa. Overall, 23 of the 112 deer RPLN samples from this preserve were found to be positive for SARS-CoV-2 RNA, with the first positive in September and the second in October 2020, and 11 of 38 deer sampled in November and all 10 deer sampled in December 2020, suggesting a rapidly increasing herd-level prevalence.

Seven counties had at least 10 samples collected, with all 11 specimens from Allamakee county being found to be SARS-CoV-2 positive, as were 21 of the 28 samples collected from Appanoose county (Table 1; Supplementary Table 1). In contrast, none of the 9 samples collected from Black Hawk county were positive, nor were the 6 RPLN samples from Henry county. While the exact reasons for this heterogeneity in PCR positive response rates are unknown, the timing of collection in relation to the SARS-CoV-2 spread in deer may play a role. For instance, the samples from Henry county were collected during April and May of 2020 during the early of the pandemic and well before the first positive sample was identified in deer. Similarly, all 9 RPLN samples tested from Black Hawk county were collected prior to the mid-November peak of reported SARS-CoV-2 cases in humans in Iowa. Together, these results suggest the widespread presence of SARS-CoV-2 RNA in deer across the State of Iowa, with strong evidence of temporal clustering.

### Whole genome based phylogeographic and phylogenomic analyses provide evidence of multiple likely reverse zoonotic spillover events of SARS-CoV-2 to deer and deer-to-deer transmission

To begin to understand the genomic diversity of SARS-CoV-2 associated with free-living and captive deer, we characterized the complete SARS-CoV-2 genomes from all 94 deer RPLN positive for the presence of viral RNA. A high-level of sequencing coverage was obtained, and Pangolin version 3.1.11 (*https://github.com/cov-lineages/pangoLEARN*, last accessed October 27, 2021) was used to identify SARS-CoV-2 lineages using previously described genome sequence analysis pipelines (13–15). Next, we used an automated vSNP pipeline (https://github.com/USDA-VS/vSNP) to identify SNPs and construct phylogenetic trees in the context of 84 additional publicly available animal origin SARS-CoV-2 isolates as well as from 372 SARS-CoV-2 isolates identified from humans in Iowa during this same period (Supplementary Table 2). All newly sequenced SARS-CoV-2 consensus genomes from deer RPLN are deposited in GISAID, and raw reads submitted to NCBI’s Short Read Archive (BioProject Number PRJNA776532).

The analysis identified a total of 12 SARS-CoV-2 lineages amongst the 94 samples from deer, with two lineages, B.1.2 (*n* = 51; 54.5%), and B.1.311 (*n* = 19; 20%) representing ~75% of all samples. Together with the next two most abundant lineages, B.1 (*n* = 7) and B.1.234 (*n* = 6), these four lineages represented ~ 88% of SARS-CoV-2 circulating amongst geographically widely distributed deer in the State (Table 2). While the number of SARS-CoV-2 sequences from Iowa is not very high (only 372 sequences are available) during this period in publicly available data, it is noteworthy that the B.1.2 was also the most abundant (~ 43.5%) SARS-CoV-2 lineage circulating in humans in Iowa (Supplementary Table 2). In contrast, the B.1.311 lineage accounting for about one fifth of the isolates in deer was relatively poorly represented (~1.6%) amongst the publicly available SARS-CoV-2 from humans in Iowa (Supplementary Table 2). Conversely, the second most abundant SARS-CoV-2 lineage in humans, B.1.565, accounting for ~ 8.1% of available sequences, was not identified amongst the sampled deer. However, given the lack of representativeness of the sampling from both humans and deer, we urge caution in overinterpreting these findings of apparent differences in prevalence of SARS-CoV-2 lineages between deer and sympatric human hosts.

**Table 2.** Recovery of SARS-CoV-2 lineages from white-tailed deer in Iowa.

| **Lineage** | **Number of**<br>**Samples** | **Proportion**<br>**(95% CI)** |
| --- | --- | --- |
| **B.1** | 7 | 7.5 (3.7-14.6) |
| **B.1.1** | 1 |  |
| **B.1.119** | 2 |  |
| **B.1.2** | 51 | 54.2 (44.2-64.0) |
| **B.1.234** | 6 | 6.4 (3.0-13.2) |
| **B.1.240** | 1 |  |
| **B.1.264** | 1 |  |
| **B.1.311** | 19 | 20.2 (13.3-29.4) |
| **B.1.362** | 2 |  |
| **B.1.400** | 2 |  |
| **B.1.459** | 1 |  |
| **B.1.596** | 1 |  |
| **Total** | 94 |  |

The temporal and geographic patterns of clustering of SARS-CoV-2 lineages, together with phylogenetic analyses, provide strong evidence of multiple likely zooanthroponotic spillovers from humans to deer. This is evidenced by the near simultaneous recovery of multiple SARS-CoV-2 lineages within temporally and geographically restricted deer herds at various locations throughout the State. For instance, the results show that a vast majority (22 of 23) of samples recovered from deer in a game preserve in Southeastern Iowa was represented by the B.1.2 lineage. The single outlier isolate was represented by lineage B.1.311 and was recovered from a deer at this preserve on November 24, 2020, in synchrony with 12 additional specimens harboring the B.1.2 lineage. Similarly, in the Yellow River State Forest area in Allamakee county, during the 5 days between December 5 and December 9, all 11 hunter-killed deer were positive for the presence of SARS-CoV-2 RNA in their RPLNs. Nine of these samples were represented by the B.1.2 lineage. The two outliers, represented by B.1.311 and B.1.459 lineages were recovered from another hunter-killed deer on the same date, December 8, 2020, as was another deer harboring the B.1.2 lineage. All these deer were hunted within a few miles of each other. A third example of near synchronous recovery of genetically distinctive SARS-CoV-2 lineages from infected deer is from the Volga River State Recreation Area in Fayette county, where 4 lineages were recovered from the RPLN of hunter-killed deer within a two-mile radius in the three days spanning December 7 through 9, 2020. A final example is from the Lake Rathbun area of Appanoose county, where on December 5, 2020, of 10 positive RPLNs harvested from hunter-killed deer, there were 5 distinctive lineages represented within a 5 mile radius – lineage B.1.311 was recovered from 6 deer RPLNs, and one each of lineages B.1.362, B.1.240, B.1.400, and B.1. Together, these findings strongly suggest multiple point sources of spillover of distinct SARS-CoV-2 lineages to captive and free-living deer.

Recent evidence suggests that experimentally infected deer readily transmit the virus to other susceptible deer between 3-5 days post infection, and the virus can be recovered from the palatine tonsils and RPLNs of infected animals for up to 21 days post-exposure (6). However, evidence of deer-to-deer transmission of the virus in free-living deer has not yet been documented. To explore evidence for potential sylvatic transmission in free-living deer, we applied a molecular epidemiologic approach to explore the temporal patterns of recovery of SARS-CoV-2 lineages from free-living deer to identify possible evidence of deer-to-deer transmission. One example is evident from the Lake Rathbun area of Appanoose county, where lineage B.1.311 was predominant amongst deer representing 14 of 21 positive samples. The first RPLN sample harboring a B.1.311 lineage was recovered on December 5, 2020 from a hunter-killed deer from this area. This was followed by additional recoveries of B.1.311 on December 8 and then again on January 2, 2021, and January 9, 2021 – more than a month apart – and with high viral loads in the lymph nodes suggestive of active infection. Together with the observation that the B.1.311 lineage was less frequently reported (based on available sequences) from humans in Iowa as well as in deer in other IA counties, the results suggest the continued circulation of this lineage amongst deer in free-living settings. However, it is important to note that in the absence of a comprehensive longitudinal study of circulating lineages, it is not possible to formally exclude the possibility that B.1.311 was circulating within humans or other hosts in this area, and the deer were repeatedly exposed to the same point source(s) of infection.

Finally, to better visualize phylogenetic relationships amongst circulating SARS-CoV-2 originating in free-living and captive deer, we generated an SNP-based maximum likelihood tree including available human and animal lineage isolates (Fig. 3). As evident from the branching patterns of the phylogram, the results highlight the presence of multiple independent but closely related SARS-CoV-2 lineages circulating amongst deer in Iowa, as well as provide strong evidence for transmission within deer as many of the genomes from individual deer shared complete genomic identity (no SNPs) or differed by between 1 and 5 SNPs. The results also highlight several branches with shared human and deer origin SARS-CoV-2 isolates circulating in Iowa that are related to but distinct from isolates previously identified from outbreaks from animals such as farmed mink or otters or other domesticated animal species. Hence, taken together, the results provide strong evidence of multiple spillover events of SARS-CoV-2 and the subsequent circulation of these strains within free-living and captive deer.

**Figure 3:**
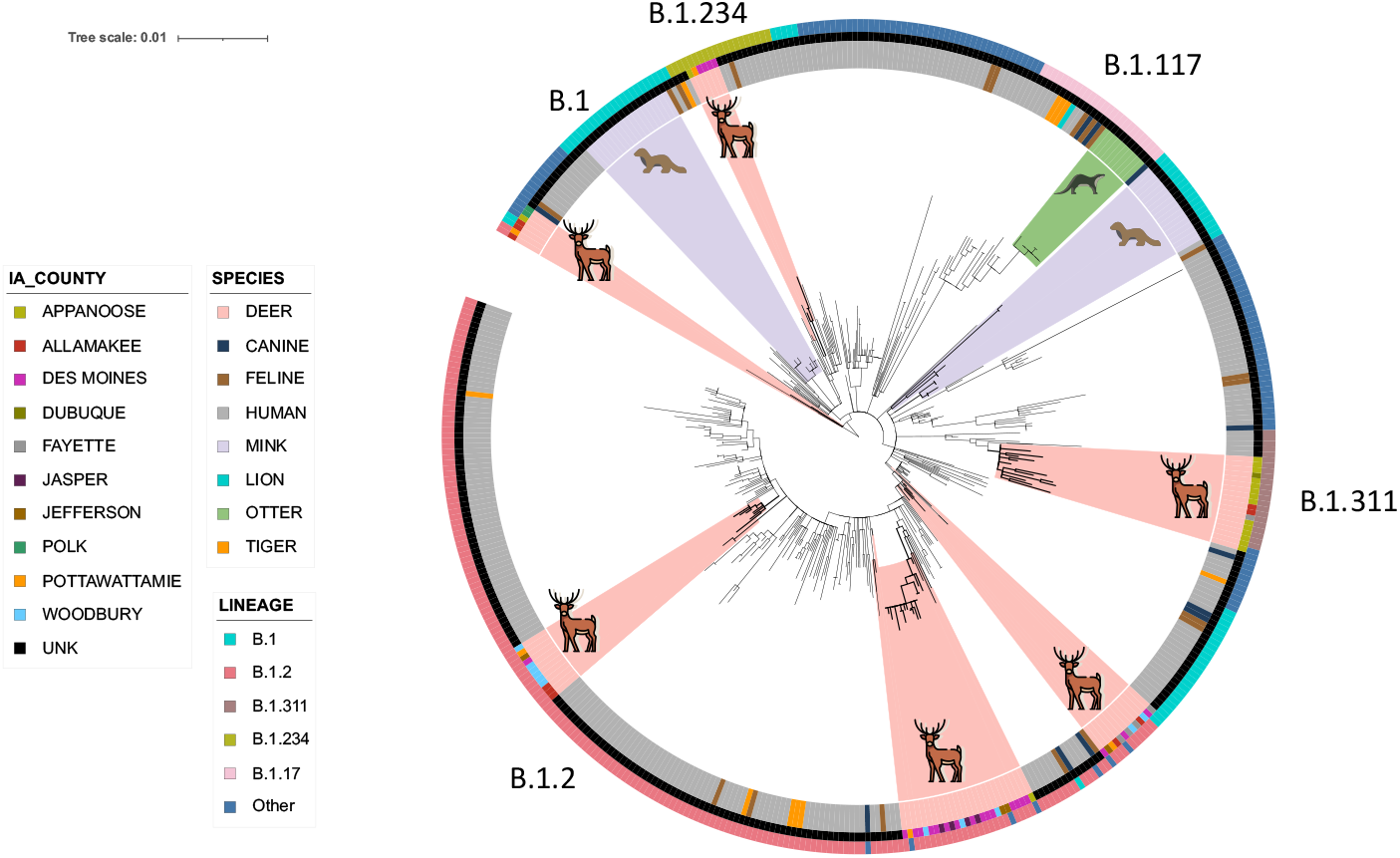
Whole genome SNP-based phylogenies of 94 SARS-CoV-2 recovered from free-living and captive white-tailed deer in Iowa. Whole genome sequences of the 94 SARS-COV-2 positive samples from deer RPLN were analyzed in the context of 92 additional publicly available animal origin SARS-CoV-2 isolates and 312 human SARS-CoV-2 isolates circulating in Iowa during this same period (Supplementary Table 2). The sequences were screened for quality thresholds, and SNP positions called against the SARS-CoV-2 reference were determined together with SNP alignments and used to assemble a maximum-likelihood phylogenetic trees using RAxML. The results show several genetically distinct clusters of animal and human SARS-CoV-2 lineages circulating within the Iowa deer herd, suggesting multiple likely spillover events from humans to deer. Several branches with shared human and deer origin SARS-CoV-2 isolates circulating in Iowa were observed. The sequences from deer were genetically distinct from isolates from previous outbreaks in farmed mink and otters but showed close clustering with SASRS-CoV-2 genomes recovered humans in Iowa.

### Broader implications for the ecology of SARS-CoV-2

Most viruses causing disease in humans have originated in animals and many are capable of transmitting between multiple host species (16, 17). The ability to infect a range of host species is a risk factor for disease emergence (18, 19). Despite this knowledge, reservoir host(s) are rarely identified and studied. Indeed, the wild animal reservoir(s) of SARS-1, SARS-CoV-2 and MERS-CoV are still not known. There have been numerous cases of isolated human-to-animal transmission of SARS-CoV-2 involving companion, farmed, and zoo animals since the COVID-19 pandemic began (8, 9, 20, 21). Our study is the first to provide evidence of widespread dissemination of SARS-CoV-2 into any free-living species, in this instance, the white-tailed deer. While the precise route(s) of transmission of SARS-Cov-2 from humans to deer are unknown, there are several ways in which deer may be exposed to the virus from humans, including through feeding in backyards or even when a susceptible deer may come in contact with potentially infectious material saliva, urine, etc.) from an infected human in forested areas or exurban environments. Deer may also become exposed to SARS-CoV-2 through contact with wastewater discharges, infected fomites, or other infected animals. Regardless of the route of transmission from humans, our results suggest that deer have the potential to emerge as a major reservoir host for SARS-CoV-2, a finding that has important implications for the future trajectory of the pandemic.

So what might be the implications of deer emerging as reservoirs of SARS-CoV-2? When pathogens infect a single host species, the population dynamics are intrinsically unstable, and an outbreak spreads rapidly through a population and then fades out as hosts either develop immunity or die from the infection. The outbreak’s trajectory depends on the basic reproductive number (R_0_) and the generation time of the infection, but this changes when a pathogen is a generalist and infects multiple host species. In this instance, the dynamics are dominated by what occurs within reservoir hosts, defined as species which can maintain the infection and from which infection is transmitted to other hosts (22).

A reservoir host can facilitate viral evolution and the emergence of lineages with increased virulence for the original host. Since many animal species already harbor an extensive array of endemic endogenous CoVs, the presence of a free-living reservoir host for SARS-CoV-2 may provide an opportunity for the virus to recombine and acquire or evolve increased fitness traits such as increased virulence, transmissibility, pathogenicity, and immune evasion (23). Evidence of some of these exists in the Denmark mink spillover event where the Y435F substitution (which conferred increased affinity of the spike protein to human ACE2) evolved after human to mink transmission (24–26). Animal reservoirs can also provide a refuge outside of a largely immune/vaccinated human population and thus increase the threat of subsequent re-emergence of ancestral genotypes into immunologically naïve or susceptible human hosts (23). An example of this is a scenario observed in the 2009 A-H1N1 (swine flu) pandemic where the virus was related both to the pandemic 1918 strain and strains circulating in the early 20^th^ century (23, 27).

Predicting how the utilization of a new host species by a virus can affect virulence in the primary host is not simple. In theory, pathogen evolution to an optimum in a single-host system is determined by a trade off with transmission, but this becomes more complicated in multiple-host systems (28). With the infection spreading so quickly through the deer population, as seen in our study, this could potentially result in fade out with insufficient susceptible deer recruits to sustain the infection within the deer population alone. Alternatively, with sizeable annual birth cohorts or invasion into areas where deer have not previously been infected, the virus may continue to spread among susceptible deer or circulate with the deer population. However, even while the dynamics in these multi-host systems can be complex, they often result in more stable dynamics with multiple reservoir hosts. The pathogens that utilize many hosts can be at a selective advantage since they are not lost during periods soon after the fade out.

Finally, the white-tailed deer is the most abundant wild cervid species in the United States, with an estimated 25 million individuals. Deer hunting is the most popular form of hunting in the United States, contributing over $20 billion to the US GDP and supporting more than 300,000 jobs in 2016 (29). Given the social relevance and economic importance of deer to the US economy, even though experimental evidence suggests that SARS-CoV-2 infected deer remain largely asymptomatic, the clinical outcomes and health implications of SARS-CoV-2 infection in free-living deer are unknown, and warrant further investigation. For these reasons, the discovery of sylvatic and enzootic transmission in a substantial fraction of free-living deer has important implications for the natural ecology and long-term persistence of the SARS-CoV-2, including through spillover to other free-living or captive animals and potential for spillback to humans.

### Study limitations

The study has several limitations: The RPLN samples tested were from only one State in the USA, and the sampling was not uniform within the State. However, while the generalizability of our findings remains to be tested, we see no reason why this scenario has also not already played out in other regions with large deer populations with opportunities for contact with humans. Another limitation of our study was that RPLN samples tested were all from 2020 and early 2021, representing the early part of the pandemic before the global dissemination of the highly successful Alpha and Delta variants. Hence, surveillance efforts with robust longitudinal sampling approaches are urgently needed to determine whether deer will become long-term reservoirs for SARS-CoV-2 and potentially assume a role as generators of novel variant viruses that may repeatedly re-emerge in humans or spillover to other animal hosts.

### Concluding comments

To help predict or prevent the emergence of the next pandemic and control infectious diseases with pandemic and panzootic potential, a better understanding of the human–animal molecular and ecological interface and its relevance to infection transmission dynamics is essential (9). Thus, we call for an urgent need to implement a more proactive and robust “One Health” approach to better understand the ecology and evolution of SARS-CoV-2 in deer and other free-living species.

## Materials and Methods

### Samples

The Iowa Department of Natural Resources (DNR) routinely collects medial retropharyngeal lymph nodes (RPLNs) from white-tailed deer across Iowa for its statewide Chronic Wasting Disease (CWD) surveillance program. Tissue samples were collected by trained field staff. Paired RPLNs were then removed and placed into separate Whirl-Paks with corresponding sample identification numbers and frozen at −20°F in a standard chest or standing freezer. A total of 283 RPLN samples collected between April 2020 to January 2021 were studied (Supplementary Table 1). An additional 60 RPLN archived samples from the 2019 deer hunting season were included as process negative control samples (Supplementary Table 1).

### RNA extraction

RPLN tissues were processed by adding 3ml UTM (Copan) to a whirl-pak bag containing the tissue and placing the bag in the stomacher on a high setting for 120 seconds. Liquid volume was recovered and centrifuged at 3,000 rpm for 5 minutes to pellet cellular debris. 400 μL of the RPLN tissue homogenate supernatant was used for viral RNA extraction with a KingFisher Flex machine (ThermoFisher Scientific) with the MagMAX Viral/Pathogen extraction kit (ThermoFisher Scientific) following the manufacturer’s instructions.

### Detection of SARS-CoV-2 viral RNA by RT-PCR

The presence of SARS-CoV-2 nucleic acid was assessed by a real-time reverse transcription-polymerase chain reaction (RT-PCR) assay using the OPTI Medical SARS-CoV-2 RT-PCR kit following the manufacturer’s instructions on an ABI 7500 Fast instrument (ThermoFisher Scientific). The OPTI Medical SARS-CoV-2 RT-PCR assay detects two different targets in the gene encoding viral nucleocapsid (N) protein coding region(12, 30). The assay is highly sensitive with a limit of detection of 0.36 copies/μl. The internal control RNase P (RP) was used to rule out human contamination. We generated a standard curve using SARS-CoV-2 RNA with a known copy number. Using the standard curve, viral RNA copies per milliliter of tissue homogenate were calculated.

To ensure assay specificity, a subset of 25 positive and 25 negative samples were additionally tested with the TaqPath kit (ThermoFisher Scientific) targeting the SARS-CoV-2 ORF1ab, N gene, and S gene (30, 31). The results were concordant with both assays. Further, to ensure samples were not inadvertently contaminated with human origin tissue or fluids during harvesting or processing, all samples were tested and found negative for the presence of human RNaseP. As a final check of assay specificity, none of the 60 RPLN samples collected in 2019 prior to the first reported case in humans in the United States were found positive for the presence of SARS-CoV-2 RNA.

### SARS-CoV-2 Genome Sequencing

Total RNAs extracted from RPLN samples was used for sequencing the whole genomes of SARS-CoV-2 as previously described (13–15). Briefly, libraries were prepared according to version 4 of the ARTIC nCoV-2019 sequencing protocol (https://artic.network/ncov-2019, last accessed October 27, 2021). We used a semi-automated workflow that employed BioMek i7 liquid handling workstations (Beckman Coulter Life Sciences, Indianapolis, IN) and MANTIS automated liquid handlers (FORMULATRIX, Bedford, MA). Short sequence reads were generated with a NovaSeq 6000 instrument (Illumina, San Diego, CA). To ensure a very high depth of coverage, the RPLN sequencing libraries were prepared in duplicate and sequenced with a SP 300 cycle reagent kit.

### SARS-CoV-2 Genome Sequence Analysis and Identification of Variants

Viral genomes were assembled with the BV-BRC SARS-Cov2 assembly service (The Bacterial and Viral Bioinformatics Resource Center, *https://www.bv-brc.org/app/comprehensivesars2analysis*, last accessed October 27, 2021, requires registration)(32). The One Codex SARS-CoV-2 variant calling and consensus assembly pipeline was used to assemble all sequences (GitHub, *https://github.com/onecodex/sars-cov-2.git*, last accessed October 27, 2021) using default parameters and a minimum read depth of three. Briefly, the pipeline uses seqtk version 1.3-r116 for sequence trimming (GitHub, *https://github.com/lh3/seqtk.git*, last accessed October 27, 2021); minimap version 2.1 (*https://github.com/lh3/minimap2*, last accessed October 27, 2021) for aligning reads against reference genome Wuhan-Hu-1 (*https://www.ncbi.nlm.nih.gov/nuccore/1798174254*; NC_045512.2)(33); samtools version 1.11 (*http://www.htslib.org*, last accessed October 27, 2021) for sequence and file manipulation(34); and iVar version 1.2.2 (*https://github.com/andersen-lab/ivar/releases*, last accessed October 29, 2021) for primer trimming and variant calling(35). Genetic lineages, VOCs, and VOIs were identified on the basis of genome sequence data and designated by Pangolin version 3.1.11 (*https://github.com/cov-lineages/pangoLEARN*, last accessed October 27, 2021) with pangoLEARN module 2021-08-024 (SARS-CoV-2 lineages, *https://cov-lineages.org/pangolin.html*, last accessed October 27, 2021). SNPs identified using vSNP (https://github.com/USDA-VS/vSNP,) SNP analysis program.

QGIS mapping software version 3.16.10 was used to portray/visualize the geographic distribution of the RPLN samples.

## Funding

The study is funded by the Huck institutes of the life sciences (SVK and VK), US Department of Agriculture National Institute of Food and Agriculture (NIFA) Award 2020-67015-32175, USFWS Wildlife and Sport Fish Restoration Program grant awards: FY20 F19AF00434 (19/20 samples), FY21 F20AF00309 (20/21 samples), and FY22 F21AF01914 (21/22 samples) and The Department of Natural Resources Fish and Game Protection Fund. This project was also partly supported by the Houston Methodist Academic Institute Infectious Diseases Fund; and supported in whole or in part with federal funds from the National Institute of Allergy and Infectious Diseases, National Institutes of Health, Department of Health and Human Services, under Contract No. 75N93019C00076 (J.J.D.)

## Author contributions

S.V.K conceptualized the study; S.V.K, V.K and R.M.R designed the project; M.S.N, M.Y, R.H.N, R.K.N, L.L, B.M.J, K.J.V, C.D.M, N.L,, K.W, A.J.K.C, R.J.O, J.J.D, J.M.M and P.J.H performed research and or analysis. All authors contributed to writing the manuscript.

## Competing interests

Authors declare that they have no competing interests.

## Data and materials availability

All SARS-CoV-2 consensus genomes are deposited in GISAID and raw reads submitted to NCBI’s Short Read Archive (BioProject Number: PRJNA776532)

## Acknowledgments

We thank Abhinay Gontu, Shubhada Chothe, Padmaja Jakka and Abirami Ravichnadran from Kuchipudi lab, Penn State for their help with tissue homogenization. Vincent Nelson, Lindsey LaBella and Corey Price from the Penn State CLIA lab for help with sample processing. We greatly acknowledge Matthew Ojeda Saavedra, Sindy Pena, Kristina Reppond, Madison N. Shyer, Jessica Cambric, Ryan Gadd, Rashi M. Thakur, Akanksha Batajoo, and Regan Mangham for their technical assistance in the genome sequencing. We are grateful to Drs Suelee Robbe-Austerman, Tod Stuber and Kristina Lantz, USDA NVSL for their invaluable assistance and wise counsel throughout this investigation.

---

**Supplementary Figure 1:**
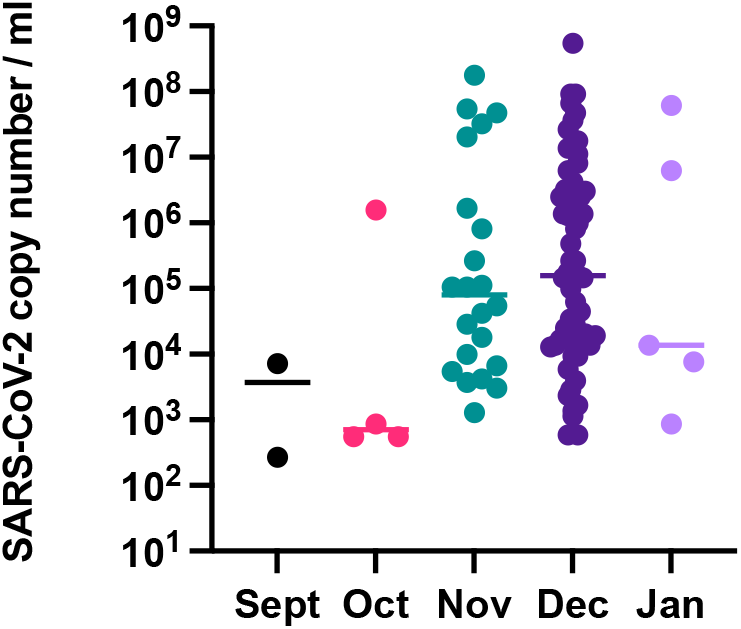
Temporal changes in distribution of SARS-CoV-2 viral genome copy numbers in white-tailed deer RPLNs. As the positivity proportion among the collected samples increased over the months of collection depicted on X axis, the viral copy numbers (y-axis) increased in a range of 268 to 5.4 × 10^8^ copies/ml with a median of 106,000 viral copies/ml.

**Supplementary Figure 2:**
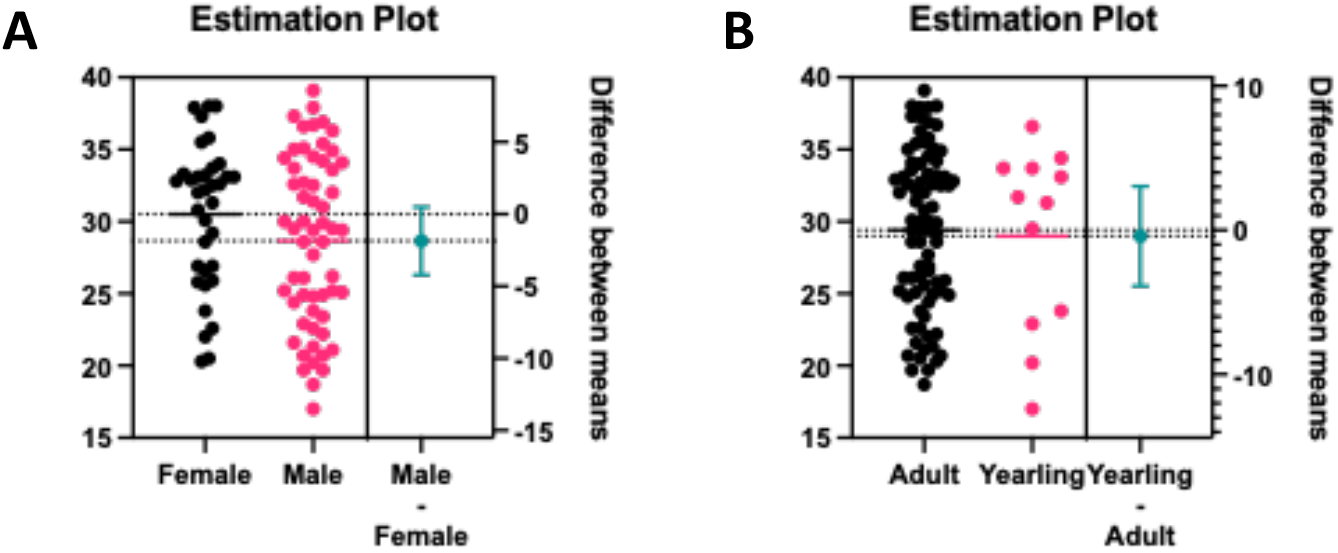
Estimation plots of gender and age associated effects on the distribution of Ct values from RPLN SARS-CoV-2 positive samples. The results show no significant difference based on Sex (A) or Age (B) in proportion of SARS-CoV-2 positive samples.

**Supplementary Table 1.** SAMPLES INCLUDED.

SAMPLES INCLUDED
| Sample ID | Date Collected | Type | Age | Sex | County | Type2 | Rt-PCR Ct value | SARSCoV-2 Line |
| --- | --- | --- | --- | --- | --- | --- | --- | --- |
| 207162 | 4/8/20 | Roadkill | Yearling | Female | Woodbury | Free living | NEG |  |
| 207159 | 4/13/20 | Roadkill | Adult | Female | Woodbury | Free living | NEG |  |
| 207160 | 4/13/20 | Roadkill | Adult | Female | Woodbury | Free living | NEG |  |
| 208000 | 4/28/20 | Preserve | Yearling | Female | SE Iowa | Captive | NEG |  |
| 208001 | 4/28/20 | Preserve | Fawn | Male | SE Iowa | Captive | NEG |  |
| 208002 | 4/28/20 | Preserve | Yearling | Female | SE Iowa | Captive | NEG |  |
| 208003 | 4/28/20 | Preserve | Adult | Female | SE Iowa | Captive | NEG |  |
| 207143 | 5/12/20 | Roadkill | Adult | Female | Woodbury | Free living | NEG |  |
| 208004 | 5/13/20 | Preserve | Adult | Male | SE Iowa | Captive | NEG |  |
| 208005 | 5/13/20 | Preserve | Adult | Female | SE Iowa | Captive | NEG |  |
| 203001 | 5/21/20 | Roadkill | Yearling | Male | Black Hawk | Free living | NEG |  |
| 207177 | 6/19/20 | Roadkill | Adult | Male | Woodbury | Free living | NEG |  |
| 207183 | 6/19/20 | Roadkill | Yearling | Female | Woodbury | Free living | NEG |  |
| 207185 | 6/19/20 | Roadkill | Adult | Female | Woodbury | Free living | NEG |  |
| 207186 | 6/19/20 | Roadkill | Adult | Female | Woodbury | Free living | NEG |  |
| 206518 | 8/19/20 | Roadkill | Adult | Female | Pottawattamie | Free living | NEG |  |
| 208300 | 8/21/20 | Preserve | Adult | Male | NE Iowa | Captive | NEG |  |
| 206519 | 9/10/20 | Roadkill | Adult | Male | Pottawattamie | Free living | NEG |  |
| 208006 | 9/18/20 | Preserve | Adult | Male | SE Iowa | Captive | NEG |  |
| 208007 | 9/18/20 | Preserve | Adult | Male | SE Iowa | Captive | NEG |  |
| 208008 | 9/18/20 | Preserve | Adult | Male | SE Iowa | Captive | NEG |  |
| 208009 | 9/18/20 | Preserve | Adult | Male | SE Iowa | Captive | NEG |  |
| 208010 | 9/18/20 | Preserve | Adult | Male | SE Iowa | Captive | NEG |  |
| 208011 | 9/18/20 | Preserve | Adult | Male | SE Iowa | Captive | NEG |  |
| 208012 | 9/18/20 | Preserve | Adult | Female | SE Iowa | Captive | NEG |  |
| 206520 | 9/20/20 | Target/Sick | Adult | Male | Pottawattamie | Free living | NEG |  |
| 207144 | 9/20/20 | Target/Sick | Adult | Female | Woodbury | Free living | NEG |  |
| 207233 | 9/21/20 | Hunter Killed | Adult | Female | Woodbury | Free living | NEG |  |
| 207234 | 9/22/20 | Hunter Killed | Adult | Female | Woodbury | Free living | NEG |  |
| 207235 | 9/22/20 | Hunter Killed | Yearling | Female | Woodbury | Free living | NEG |  |
| 207236 | 9/22/20 | Hunter Killed | Adult | Female | Woodbury | Free living | NEG |  |
| 207237 | 9/22/20 | Hunter Killed | Adult | Female | Woodbury | Free living | NEG |  |
| 207238 | 9/22/20 | Hunter Killed | Adult | Female | Woodbury | Free living | NEG |  |
| 206521 | 9/23/20 | Roadkill | Adult | Female | Pottawattamie | Free living | NEG |  |
| 206522 | 9/24/20 | Roadkill | Adult | Female | Pottawattamie | Free living | NEG |  |
| 206523 | 9/24/20 | Roadkill | Adult | Male | Pottawattamie | Free living | NEG |  |
| 203033 | 9/25/20 | Hunter Killed | Adult | Female | Dubuque | Free living | NEG |  |
| 203034 | 9/25/20 | Hunter Killed | Yearling | Female | Dubuque | Free living | NEG |  |

**SAMPLES INCLUDED**
| Sample ID | Date Collected | Type | Age | Sex | County | Type2 | Rt-PCR Ct value | SARSCoV-2 Line |
| --- | --- | --- | --- | --- | --- | --- | --- | --- |
| 207239 | 9/26/20 | Hunter Killed | Adult | Female | Woodbury | Free living | NEG |  |
| 207240 | 9/26/20 | Hunter Killed | Adult | Female | Woodbury | Free living | NEG |  |
| 208013 | 9/28/20 | Preserve | Adult | Male | SE Iowa | Captive | NEG |  |
| 208014 | 9/28/20 | Preserve | Adult | Female | SE Iowa | Captive | NEG |  |
| 208015 | 9/28/20 | Preserve | Adult | Male | SE Iowa | Captive | NEG |  |
| 208016 | 9/28/20 | Preserve | Adult | Male | SE Iowa | Captive | NEG |  |
| 208017 | 9/28/20 | Preserve | Adult | Male | SE Iowa | Captive | 39.1 | B.1.2 |
| 208018 | 9/28/20 | Preserve | Adult | Male | SE Iowa | Captive | NEG |  |
| 208019 | 9/28/20 | Preserve | Adult | Male | SE Iowa | Captive | NEG |  |
| 208020 | 9/28/20 | Preserve | Adult | Male | SE Iowa | Captive | NEG |  |
| 208021 | 9/28/20 | Preserve | Adult | Male | SE Iowa | Captive | NEG |  |
| 208022 | 9/28/20 | Preserve | Adult | Male | SE Iowa | Captive | NEG |  |
| 208023 | 9/28/20 | Preserve | Adult | Male | SE Iowa | Captive | NEG |  |
| 208024 | 9/28/20 | Preserve | Adult | Male | SE Iowa | Captive | NEG |  |
| 208025 | 9/28/20 | Preserve | Adult | Male | SE Iowa | Captive | NEG |  |
| 207241 | 9/29/20 | Hunter Killed | Adult | Female | Woodbury | Free living | NEG |  |
| 207242 | 9/29/20 | Hunter Killed | Adult | Female | Woodbury | Free living | NEG |  |
| 207337 | 9/30/20 | Roadkill | Adult | Male | Woodbury | Free living | 34.1 | B.1.2 |
| 207243 | 10/5/20 | Hunter Killed | Adult | Female | Woodbury | Free living | NEG |  |
| 207244 | 10/5/20 | Hunter Killed | Adult | Female | Woodbury | Free living | NEG |  |
| 208026 | 10/8/20 | Preserve | Adult | Male | SE Iowa | Captive | NEG |  |
| 208027 | 10/8/20 | Preserve | Adult | Male | SE Iowa | Captive | NEG |  |
| 208028 | 10/8/20 | Preserve | Adult | Male | SE Iowa | Captive | NEG |  |
| 208029 | 10/8/20 | Preserve | Adult | Male | SE Iowa | Captive | NEG |  |
| 208030 | 10/8/20 | Preserve | Adult | Male | SE Iowa | Captive | NEG |  |
| 208031 | 10/8/20 | Preserve | Adult | Male | SE Iowa | Captive | NEG |  |
| 208032 | 10/8/20 | Preserve | Adult | Male | SE Iowa | Captive | NEG |  |
| 208033 | 10/8/20 | Preserve | Adult | Male | SE Iowa | Captive | NEG |  |
| 208034 | 10/8/20 | Preserve | Adult | Male | SE Iowa | Captive | NEG |  |
| 208035 | 10/8/20 | Preserve | Adult | Male | SE Iowa | Captive | NEG |  |
| 208036 | 10/8/20 | Preserve | Adult | Male | SE Iowa | Captive | NEG |  |
| 208037 | 10/8/20 | Preserve | Adult | Male | SE Iowa | Captive | NEG |  |
| 208038 | 10/8/20 | Preserve | Adult | Male | SE Iowa | Captive | NEG |  |
| 208039 | 10/8/20 | Preserve | Adult | Male | SE Iowa | Captive | NEG |  |
| 208040 | 10/8/20 | Preserve | Adult | Male | SE Iowa | Captive | NEG |  |
| 208041 | 10/8/20 | Preserve | Adult | Male | SE Iowa | Captive | NEG |  |
| 208042 | 10/8/20 | Preserve | Adult | Male | SE Iowa | Captive | NEG |  |
| 208043 | 10/8/20 | Preserve | Adult | Female | SE Iowa | Captive | 38 | B.1.2 |
| 207245 | 10/9/20 | Hunter Killed | Adult | Female | Woodbury | Free living | NEG |  |

**SAMPLES INCLUDED**
| Sample ID | Date Collected | Type | Age | Sex | County | Type2 | Rt-PCR Ct value | SARSCoV-2 Line |
| --- | --- | --- | --- | --- | --- | --- | --- | --- |
| 207349 | 10/11/20 | Roadkill | Adult | Female | Woodbury | Free living | NEG |  |
| 206524 | 10/12/20 | Roadkill | Adult | Female | Pottawattamie | Free living | NEG |  |
| 202276 | 10/13/20 | Hunter Killed | Adult | Female | Black Hawk | Free living | NEG |  |
| 208044 | 10/13/20 | Preserve | Adult | Female | SE Iowa | Captive | NEG |  |
| 208045 | 10/13/20 | Preserve | Adult | Male | SE Iowa | Captive | NEG |  |
| 208046 | 10/13/20 | Preserve | Adult | Male | SE Iowa | Captive | NEG |  |
| 208047 | 10/13/20 | Preserve | Adult | Male | SE Iowa | Captive | NEG |  |
| 208048 | 10/13/20 | Preserve | Adult | Male | SE Iowa | Captive | NEG |  |
| 208049 | 10/13/20 | Preserve | Adult | Male | SE Iowa | Captive | NEG |  |
| 208050 | 10/13/20 | Preserve | Adult | Male | SE Iowa | Captive | NEG |  |
| 208051 | 10/13/20 | Preserve | Adult | Male | SE Iowa | Captive | NEG |  |
| 208052 | 10/13/20 | Preserve | Adult | Male | SE Iowa | Captive | NEG |  |
| 206525 | 10/13/20 | Roadkill | Adult | Male | Pottawattamie | Free living | NEG |  |
| 207246 | 10/13/20 | Target/Sick | Adult | Female | Woodbury | Free living | NEG |  |
| 202277 | 10/14/20 | Hunter Killed | Yearling | Female | Black Hawk | Free living | NEG |  |
| 206526 | 10/14/20 | Target/Sick | Adult | Female | Pottawattamie | Free living | NEG |  |
| 206527 | 10/15/20 | Roadkill | Adult | Female | Pottawattamie | Free living | 38 | B.1.2 |
| 207385 | 10/15/20 | Roadkill | Adult | Female | Woodbury | Free living | NEG |  |
| 202278 | 10/16/20 | Hunter Killed | Yearling | Female | Black Hawk | Free living | NEG |  |
| 202275 | 10/17/20 | Hunter Killed | Yearling | Female | Black Hawk | Free living | NEG |  |
| 208325 | 10/17/20 | Target/Escape | Adult | Male | Washington | Captive | NEG |  |
| 208383 | 10/19/20 | Target/Escape | Adult | Male | Keokuk | Captive | NEG |  |
| 207247 | 10/21/20 | Hunter Killed | Adult | Female | Woodbury | Free living | NEG |  |
| 207248 | 10/21/20 | Roadkill | Adult | Female | Woodbury | Free living | NEG |  |
| 208053 | 10/22/20 | Preserve | Adult | Male | SE Iowa | Captive | NEG |  |
| 208054 | 10/22/20 | Preserve | Adult | Male | SE Iowa | Captive | NEG |  |
| 208055 | 10/22/20 | Preserve | Adult | Male | SE Iowa | Captive | NEG |  |
| 208056 | 10/22/20 | Preserve | Adult | Male | SE Iowa | Captive | NEG |  |
| 208057 | 10/22/20 | Preserve | Adult | Male | SE Iowa | Captive | NEG |  |
| 208058 | 10/22/20 | Preserve | Adult | Male | SE Iowa | Captive | NEG |  |
| 208059 | 10/22/20 | Preserve | Adult | Male | SE Iowa | Captive | NEG |  |
| 208060 | 10/22/20 | Preserve | Adult | Male | SE Iowa | Captive | NEG |  |
| 208061 | 10/22/20 | Preserve | Adult | Male | SE Iowa | Captive | NEG |  |
| 208062 | 10/22/20 | Preserve | Adult | Male | SE Iowa | Captive | NEG |  |
| 208063 | 10/22/20 | Preserve | Adult | Male | SE Iowa | Captive | NEG |  |
| 208064 | 10/22/20 | Preserve | Adult | Male | SE Iowa | Captive | NEG |  |
| 208065 | 10/22/20 | Preserve | Adult | Male | SE Iowa | Captive | NEG |  |
| 208066 | 10/22/20 | Preserve | Adult | Male | SE Iowa | Captive | NEG |  |
| 208068 | 10/22/20 | Preserve | Adult | Male | SE Iowa | Captive | NEG |  |
| 200334 | 10/27/20 | Hunter Killed | Adult | Male | Appanoose | Free living | NEG |  |

**SAMPLES INCLUDED**
| Sample ID | Date Collected | Type | Age | Sex | County | Type2 | Rt-PCR Ct value | SARSCoV-2 Line |
| --- | --- | --- | --- | --- | --- | --- | --- | --- |
| 206528 | 10/27/20 | Roadkill | Adult | Female | Pottawattamie | Free living | NEG |  |
| 208326 | 10/27/20 | Target/Escape | Adult | Female | Van Buren | Captive | NEG |  |
| 207219 | 10/27/20 | Roadkill | Yearling | Female | Woodbury | Free living | NEG |  |
| 207249 | 10/27/20 | Hunter Killed | Adult | Female | Woodbury | Free living | NEG |  |
| 207431 | 10/28/20 | Roadkill | Adult | Female | Woodbury | Free living | 25.9 | B.1.2 |
| 206529 | 10/29/20 | Roadkill | Adult | Male | Pottawattamie | Free living | NEG |  |
| 207149 | 10/29/20 | Hunter Killed | Adult | Male | Woodbury | Free living | NEG |  |
| 200284 | 10/30/20 | Hunter Killed | Yearling | Female | Appanoose | Free living | NEG |  |
| 207437 | 10/31/20 | Roadkill | Adult | Male | Woodbury | Free living | 37.3 | B.1.2 |
| 200340 | 11/2/20 November | Hunter Killed | Adult | Male | Appanoose | Free living | NEG |  |
| 207250 | 11/2/20 November | Hunter Killed | Adult | Female | Woodbury | Free living | NEG |  |
| 208069 | 11/3/20 November | Preserve | Adult | Male | SE Iowa | Captive | NEG |  |
| 208070 | 11/3/20 November | Preserve | Adult | Male | SE Iowa | Captive | NEG |  |
| 208071 | 11/3/20 November | Preserve | Adult | Male | SE Iowa | Captive | NEG |  |
| 208072 | 11/3/20 November | Preserve | Adult | Male | SE Iowa | Captive | NEG |  |
| 208073 | 11/3/20 November | Preserve | Adult | Male | SE Iowa | Captive | NEG |  |
| 208074 | 11/3/20 November | Preserve | Adult | Male | SE Iowa | Captive | NEG |  |
| 208075 | 11/3/20 November | Preserve | Adult | Male | SE Iowa | Captive | NEG |  |
| 208076 | 11/3/20 November | Preserve | Adult | Male | SE Iowa | Captive | NEG |  |
| 208077 | 11/3/20 November | Preserve | Adult | Male | SE Iowa | Captive | NEG |  |
| 208078 | 11/3/20 November | Preserve | Adult | Male | SE Iowa | Captive | NEG |  |
| 208079 | 11/3/20 November | Preserve | Adult | Male | SE Iowa | Captive | NEG |  |
| 208080 | 11/3/20 November | Preserve | Adult | Male | SE Iowa | Captive | NEG |  |
| 208081 | 11/3/20 November | Preserve | Adult | Male | SE Iowa | Captive | NEG |  |
| 208082 | 11/3/20 November | Preserve | Adult | Female | SE Iowa | Captive | NEG |  |
| 208083 | 11/3/20 November | Preserve | Adult | Female | SE Iowa | Captive | NEG |  |
| 208084 | 11/3/20 November | Preserve | Adult | Male | SE Iowa | Captive | NEG |  |
| 206530 | 11/3/20 November | Hunter Killed | Adult | Male | Pottawattamie | Free living | NEG |  |
| 207471 | 11/3/20 November | Roadkill | Yearling | Male | Woodbury | Free living | NEG |  |
| 203062 | 11/4/20 November | Hunter Killed | Adult | Female | Jackson | Free living | NEG |  |
| 203063 | 11/4/20 November | Hunter Killed | Adult | Female | Jackson | Free living | NEG |  |
| 203064 | 11/4/20 November | Hunter Killed | Adult | Female | Jackson | Free living | NEG |  |
| 206531 | 11/4/20 November | Hunter Killed | Adult | Female | Pottawattamie | Free living | NEG |  |
| 203132 | 11/5/20 November | Hunter Killed | Yearling | Female | Dubuque | Free living | NEG |  |
| 203134 | 11/5/20 November | Hunter Killed | Yearling | Female | Dubuque | Free living | NEG |  |
| 203136 | 11/5/20 November | Roadkill | Adult | Male | Dubuque | Free living | NEG |  |
| 203140 | 11/5/20 November | Hunter Killed | Adult | Female | Dubuque | Free living | NEG |  |
| 206532 | 11/5/20 November | Roadkill | Adult | Female | Pottawattamie | Free living | NEG |  |
| 200349 | 11/6/20 November | Roadkill | Yearling | Male | Appanoose | Free living | NEG |  |
| 206407 | 11/6/20 November | Hunter Killed | Adult | Male | Pottawattamie | Free living | 18.7 | B.1.2 |

**SAMPLES INCLUDED**
| Sample ID | Date Collected | Type | Age | Sex | County | Type2 | Rt-PCR Ct value | SARSCoV-2 Line |
| --- | --- | --- | --- | --- | --- | --- | --- | --- |
| 207524 | 11/6/20 November | Roadkill | Adult | Female | Woodbury | Free living | NEG |  |
| 207530 | 11/7/20 November | Roadkill | Adult | Female | Woodbury | Free living | 22 | B.1.2 |
| 207531 | 11/8/20 November | Roadkill | Adult | Female | Woodbury | Free living | 25.8 | B.1.2 |
| 203193 | 11/10/20 | Hunter Killed | Adult | Male | Dubuque | Free living | NEG |  |
| 203189 | 11/12/20 | Hunter Killed | Adult | Male | Dubuque | Free living | NEG |  |
| 206670 | 11/12/20 | Roadkill | Yearling | Male | Pottawattamie | Free living | NEG |  |
| 202282 | 11/13/20 | Hunter Killed | Adult | Female | Black Hawk | Free living | NEG |  |
| 202283 | 11/13/20 | Hunter Killed | Adult | Female | Black Hawk | Free living | NEG |  |
| 202284 | 11/13/20 | Roadkill | Adult | Female | Black Hawk | Free living | NEG |  |
| 202985 | 11/13/20 | Hunter Killed | Adult | Male | Black Hawk | Free living | NEG |  |
| 208085 | 11/13/20 | Preserve | Adult | Male | SE Iowa | Captive | NEG |  |
| 208086 | 11/13/20 | Preserve | Adult | Male | SE Iowa | Captive | NEG |  |
| 208087 | 11/13/20 | Preserve | Adult | Male | SE Iowa | Captive | NEG |  |
| 208088 | 11/13/20 | Preserve | Adult | Male | SE Iowa | Captive | NEG |  |
| 208089 | 11/13/20 | Preserve | Adult | Male | SE Iowa | Captive | NEG |  |
| 208090 | 11/13/20 | Preserve | Adult | Male | SE Iowa | Captive | NEG |  |
| 208091 | 11/13/20 | Preserve | Adult | Male | SE Iowa | Captive | NEG |  |
| 208092 | 11/13/20 | Preserve | Adult | Male | SE Iowa | Captive | NEG |  |
| 208093 | 11/13/20 | Preserve | Adult | Male | SE Iowa | Captive | NEG |  |
| 208094 | 11/13/20 | Preserve | Adult | Male | SE Iowa | Captive | NEG |  |
| 208327 | 11/13/20 | Preserve | Adult | Male | SE Iowa | Captive | NEG |  |
| 207150 | 11/15/20 | Roadkill | Adult | Male | Woodbury | Free living | 31 | B.1.2 |
| 208328 | 11/17/20 | Preserve | Adult | Male | SE Iowa | Captive | NEG |  |
| 208329 | 11/17/20 | Preserve | Adult | Male | SE Iowa | Captive | 34.5 | B.1.2 |
| 208330 | 11/17/20 | Preserve | Adult | Male | SE Iowa | Captive | 35.1 | B.1.2 |
| 207579 | 11/17/20 | Roadkill | Adult | Female | Woodbury | Free living | 28.6 | B.1 |
| 208331 | 11/18/20 | Preserve | Adult | Male | SE Iowa | Captive | 36.7 | B.1.1 |
| 200030 | 11/19/20 | Target/Escape | Adult | Male | Polk | Captive | NEG |  |
| 208067 | 11/20/20 | Preserve | Adult | Male | Des Moines | Captive | NEG |  |
| 203253 | 11/22/20 | Target/Sick | Adult | Male | Dubuque | Free living | NEG |  |
| 208095 | 11/24/20 | Preserve | Adult | Male | SE Iowa | Captive | 34.9 | B.1.2 |
| 208096 | 11/24/20 | Preserve | Adult | Male | SE Iowa | Captive | NEG |  |
| 208097 | 11/24/20 | Preserve | Adult | Male | SE Iowa | Captive | NEG |  |
| 208098 | 11/24/20 | Preserve | Adult | Male | SE Iowa | Captive | 34.2 | B.1.2 |
| 208099 | 11/24/20 | Preserve | Adult | Male | SE Iowa | Captive | 32.7 | B.1.2 |
| 208100 | 11/24/20 | Preserve | Adult | Male | SE Iowa | Captive | 33.6 | B.1.2 |
| 208101 | 11/24/20 | Preserve | Adult | Male | SE Iowa | Captive | 35.4 | B.1.2 |
| 208102 | 11/24/20 | Preserve | Adult | Male | SE Iowa | Captive | 20.7 | B.1.2 |
| 208103 | 11/24/20 | Preserve | Adult | Male | SE Iowa | Captive | 31.4 | B.1.2 |
| 208104 | 11/24/20 | Preserve | Adult | Male | SE Iowa | Captive | 29.9 | B.1.2 |

**SAMPLES INCLUDED**
| Sample ID | Date Collected | Type | Age | Sex | County | Type2 | Rt-PCR Ct value | SARSCoV-2 Line |
| --- | --- | --- | --- | --- | --- | --- | --- | --- |
| 208105 | 11/24/20 | Preserve | Adult | Male | SE Iowa | Captive | 32 | B.1.311 |
| 208106 | 11/24/20 | Preserve | Adult | Male | SE Iowa | Captive | 30 | B.1.2 |
| 208107 | 11/24/20 | Preserve | Adult | Male | SE Iowa | Captive | 21.3 | B.1.2 |
| 207589 | 11/24/20 | Hunter Killed | Adult | Female | Woodbury | Free living | 20.5 | B.1.2 |
| 207391 | 11/29/20 | Target/Sick | Adult | Male | Woodbury | Free living | 30 | B.1.2 |
| 207253 | 11/30/20 | Hunter Killed | Adult | Female | Woodbury | Free living | 26.9 | B.1.311 |
| 207770 | 12/2/20 December | Roadkill | Adult | Female | Woodbury | Free living | 32.8 | B.1.311 |
| 203164 | 12/3/20 December | Hunter Killed | Adult | Male | Dubuque | Free living | NEG |  |
| 203229 | 12/4/20 December | Hunter Killed | Adult | Male | Dubuque | Free living | NEG |  |
| 204472 | 12/5/20 December | Hunter Killed | Adult | Male | Allamakee | Free living | 28.6 | B.1.2 |
| 204473 | 12/5/20 December | Hunter Killed | Adult | Female | Allamakee | Free living | 29.2 | B.1.2 |
| 204478 | 12/5/20 December | Hunter Killed | Adult | Female | Allamakee | Free living | 33.1 | B.1.2 |
| 200276 | 12/5/20 December | Hunter Killed | Adult | Female | Appanoose | Free living | NEG |  |
| 200277 | 12/5/20 December | Hunter Killed | Adult | Female | Appanoose | Free living | NEG |  |
| 200566 | 12/5/20 December | Hunter Killed | Adult | Male | Appanoose | Free living | 19.7 | B.1.311 |
| 200567 | 12/5/20 December | Hunter Killed | Adult | Female | Appanoose | Free living | 32.6 | B.1.311 |
| 200568 | 12/5/20 December | Hunter Killed | Adult | Male | Appanoose | Free living | 26.1 | B.1.311 |
| 200569 | 12/5/20 December | Hunter Killed | Adult | Female | Appanoose | Free living | 30.8 | B.1.311 |
| 200570 | 12/5/20 December | Hunter Killed | Adult | Male | Appanoose | Free living | 19.7 | B.1.311 |
| 200571 | 12/5/20 December | Hunter Killed | Adult | Male | Appanoose | Free living | 22.2 | B.1.362 |
| 200572 | 12/5/20 December | Hunter Killed | Adult | Male | Appanoose | Free living | 25.1 | B.1.240 |
| 200573 | 12/5/20 December | Hunter Killed | Adult | Female | Appanoose | Free living | 32 | B.1.311 |
| 200574 | 12/5/20 December | Hunter Killed | Adult | Male | Appanoose | Free living | 29.4 | B.1.400 |
| 200772 | 12/5/20 December | Hunter Killed | Adult | Male | Appanoose | Free living | 26.1 | B.1 |
| 203239 | 12/5/20 December | Hunter Killed | Adult | Male | Dubuque | Free living | 32.6 | B.1.2 |
| 207649 | 12/5/20 December | Hunter Killed | Adult | Male | Woodbury | Free living | 28.6 | B.1.596 |
| 201788 | 12/6/20 December | Hunter Killed | Yearling | Male | Polk | Free living | 33.7 | B.1.234 |
| 201793 | 12/6/20 December | Hunter Killed | Adult | Female | Polk | Free living | 30.1 | B.1.119 |
| 201794 | 12/6/20 December | Hunter Killed | Yearling | Male | Polk | Free living | 36.6 | B.1.234 |
| 201795 | 12/6/20 December | Hunter Killed | Adult | Female | Polk | Free living | 33.2 | B.1.234 |
| 201796 | 12/6/20 December | Hunter Killed | Yearling | Female | Polk | Free living | 33.7 | B.1.264 |
| 201797 | 12/6/20 December | Hunter Killed | Yearling | Male | Polk | Free living | 34.4 | B.1.400 |
| 205795 | 12/7/20 December | Hunter Killed | Adult | Female | Allamakee | Free living | 33.3 | B.1.2 |
| 204363 | 12/7/20 December | Hunter Killed | Adult | Male | Fayette | Free living | 37.9 | B.1 |
| 204364 | 12/7/20 December | Hunter Killed | Adult | Male | Fayette | Free living | 36.9 | B.1.2 |
| 204374 | 12/7/20 December | Hunter Killed | Yearling | Female | Fayette | Free living | NEG |  |
| 205750 | 12/7/20 December | Hunter Killed | Adult | Female | Fayette | Free living | 37.9 | B.1.234 |
| 205751 | 12/7/20 December | Hunter Killed | Yearling | Male | Fayette | Free living | NEG |  |
| 205752 | 12/7/20 December | Hunter Killed | Yearling | Male | Fayette | Free living | NEG |  |
| 203522 | 12/8/20 December | Hunter Killed | Adult | Female | Allamakee | Free living | 35.8 | B.1.2 |

**SAMPLES INCLUDED**
| Sample ID | Date Collected | Type | Age | Sex | County | Type2 | Rt-PCR Ct value | SARSCoV-2 Line |
| --- | --- | --- | --- | --- | --- | --- | --- | --- |
| 203525 | 12/8/20 December | Hunter Killed | Yearling | Male | Allamakee | Free living | 20.2 | B.1.311 |
| 203526 | 12/8/20 December | Hunter Killed | Adult | Female | Allamakee | Free living | 26.6 | B.1.459 |
| 200191 | 12/8/20 December | Hunter Killed | Adult | Male | Appanoose | Free living | 25.2 | B.1.311 |
| 200192 | 12/8/20 December | Hunter Killed | Adult | Male | Appanoose | Free living | 35 | B.1.311 |
| 200193 | 12/8/20 December | Hunter Killed | Adult | Male | Appanoose | Free living | 24.9 | B.1.311 |
| 200194 | 12/8/20 December | Hunter Killed | Adult | Female | Appanoose | Free living | 25.6 | B.1.311 |
| 200195 | 12/8/20 December | Hunter Killed | Adult | Female | Appanoose | Free living | 33.1 | B.1.311 |
| 200459 | 12/8/20 December | Hunter Killed | Adult | Male | Appanoose | Free living | NEG |  |
| 205769 | 12/8/20 December | Hunter Killed | Adult | Male | Fayette | Free living | 29.4 | B.1.2 |
| 203700 | 12/9/20 December | Hunter Killed | Adult | Male | Allamakee | Free living | 23.4 | B.1.2 |
| 203701 | 12/9/20 December | Hunter Killed | Adult | Male | Allamakee | Free living | 20.7 | B.1.2 |
| 203704 | 12/9/20 December | Hunter Killed | Yearling | Male | Allamakee | Free living | 29.5 | B.1.2 |
| 203705 | 12/9/20 December | Hunter Killed | Yearling | Female | Allamakee | Free living | 31.3 | B.1.2 |
| 201020 | 12/9/20 December | Hunter Killed | Adult | Male | Dickinson | Free living | NEG |  |
| 204668 | 12/9/20 December | Hunter Killed | Adult | Female | Fayette | Free living | 32.5 | B.1.311 |
| 204669 | 12/9/20 December | Hunter Killed | Adult | Female | Fayette | Free living | 32.2 | B.1.234 |
| 208108 | 12/10/20 | Preserve | Adult | Male | SE Iowa | Captive | 32.5 | B.1.2 |
| 208109 | 12/10/20 | Preserve | Adult | Male | SE Iowa | Captive | 26.2 | B.1.2 |
| 208110 | 12/10/20 | Preserve | Adult | Male | SE Iowa | Captive | 24.9 | B.1.2 |
| 208111 | 12/10/20 | Preserve | Adult | Male | SE Iowa | Captive | 24.4 | B.1.2 |
| 208112 | 12/10/20 | Preserve | Adult | Male | SE Iowa | Captive | 27.7 | B.1.2 |
| 208113 | 12/10/20 | Preserve | Adult | Male | SE Iowa | Captive | 25.2 | B.1.2 |
| 208114 | 12/10/20 | Preserve | Adult | Male | SE Iowa | Captive | 21.1 | B.1.2 |
| 208115 | 12/10/20 | Preserve | Adult | Male | SE Iowa | Captive | 24.8 | B.1.2 |
| 208116 | 12/10/20 | Preserve | Adult | Male | SE Iowa | Captive | 22.6 | B.1.2 |
| 201820 | 12/10/20 | Hunter Killed | Yearling | Male | Jasper | Free living | 17 | B.1 |
| 201821 | 12/10/20 | Hunter Killed | Adult | Female | Jasper | Free living | 22.6 | B.1 |
| 201833 | 12/10/20 | Hunter Killed | Yearling | Male | Jasper | Free living | 23.8 | B.1 |
| 201835 | 12/10/20 | Hunter Killed | Adult | Female | Jasper | Free living | 26.9 | B.1 |
| 201741 | 12/10/20 | Roadkill | Adult | Male | Webster | Free living | NEG |  |
| 207707 | 12/10/20 | Roadkill | Yearling | Male | Woodbury | Free living | 31.7 | B.1.2 |
| 200069 | 12/15/20 | Hunter Killed | Adult | Female | Appanoose | Free living | 32.9 | B.1.119 |
| 206739 | 12/15/20 | Roadkill | Yearling | Male | Pottawattamie | Free living | 22.9 | B.1.2 |
| 208334 | 12/17/20 | Preserve | Adult | Male | SE Iowa | Captive | 36.3 | B.1.2 |
| 208335 | 12/17/20 | Preserve | Adult | Male | SE Iowa | Captive | NEG |  |
| 208336 | 12/17/20 | Preserve | Adult | Male | SE Iowa | Captive | NEG |  |
| 206533 | 12/18/20 | Roadkill | Adult | Female | Pottawattamie | Free living | NEG |  |
| 206534 | 12/18/20 | Roadkill | Adult | Unknown | Pottawattamie | Free living | NEG |  |
| 206535 | 12/22/20 | Roadkill | Adult | Male | Pottawattamie | Free living | 21.6 | B.1.2 |
| 208117 | 12/23/20 | Preserve | Adult | Male | SE Iowa | Captive | 29.5 | B.1.2 |

**SAMPLES INCLUDED**
| Sample_ID | Date Collected | Type | Age | Sex | County | Type2 | Rt-PCR Ct value | SARSCoV-2 Line |
| --- | --- | --- | --- | --- | --- | --- | --- | --- |
| 206978 | 12/28/20 | Roadkill | Adult | Female | Pottawattamie | Free living | 35.5 | B.1.2 |
| 200442 | 1/9/21 | Hunter Killed | Adult | Female | Appanoose | Free living | 37.3 | B.1.311 |
| 200443 | 1/9/21 | Hunter Killed | Adult | Female | Appanoose | Free living | 23.8 | B.1.311 |
| 200444 | 1/9/21 | Hunter Killed | Yearling | Female | Appanoose | Free living | 33.1 | B.1.311 |
| 200862 | 1/9/21 | Hunter Killed | Adult | Female | Appanoose | Free living | 20.3 | B.1.362 |
| 200866 | 1/9/21 | Hunter Killed | Adult | Female | Appanoose | Free living | 34 | B.1.234 |

**SAMPLES EXCLUDED**
| Sample_ID | Date_Collected | Sample | Age | Sex | County | Location | Location2 | Comment |
| --- | --- | --- | --- | --- | --- | --- | --- | --- |
| 208332 | 11/19/20 | Preserve | Adult | Female | SE Iowa | Captive_ELK | Preserve 3 | Wrong Species |
| 208333 | 11/19/20 | Preserve | Adult | Female | SE Iowa | Captive_ELK | Preserve 3 | Wrong Species |
| 200934 | 2/2/21 | Target/Escape | Adult | Male | Monroe | Captive | Found_dead | ?? Date / Location |
| 208301 | Not specified | Preserve | Adult | Unknown | NE Iowa | Captive | Preserve 1 | Missing Date |
| 208302 | Not specified | Preserve | Adult | Unknown | NE Iowa | Captive | Preserve 1 | Missing Date |
| 208303 | Not specified | Preserve | Adult | Unknown | NE Iowa | Captive | Preserve 1 | Missing Date |

**PRE-PANDEMIC SAMPLES INCLUDED**
| Sample_ID | Date_Collected | Sample | Age | Sex | County | Location | RT-PCR results | Column1 |
| --- | --- | --- | --- | --- | --- | --- | --- | --- |
| 193076 | 43562 | Roadkill | Yearling | Male | Clinton | Rural | NEG |  |
| 193057 | 43726 | Roadkill | Yearling | Male | Black Hawk | Rural | NEG |  |
| 197229 | 43752 | Hunter Killed | Adult | Male | Crawford | Rural | NEG |  |
| 193146 | 43754 | Target/Sick | Adult | Female | Clinton | Rural | NEG |  |
| 193148 | 43754 | Hunter Killed | Adult | F | Clinton | Public_land | NEG |  |
| 193218 | 43781 | Hunter Killed | Adult | Male | Clinton | Public_land | NEG |  |
| 193293 | 43783 | Hunter Killed | Adult | Female | Scott | Rural | NEG |  |
| 193298 | 43784 | Hunter Killed | Adult | Male | Johnson | Rural | NEG |  |
| 193304 | 43784 | Target/Sick | Yearling | Male | Scott | Public_land | NEG |  |
| 194231 | 43784 | Target/Sick | Adult | Male | Black Hawk | Public_land | NEG |  |
| 198715 | 43785 | Hunter Killed | Adult | Male | Warren | Rural | NEG |  |
| 193305 | 43787 | Hunter Killed | Adult | Female | Scott | Public_land | NEG |  |
| 193306 | 43787 | Hunter Killed | Adult | Female | Scott | Public_land | NEG |  |
| 194229 | 43787 | Roadkill | Yearling | Female | Black Hawk | Rural | NEG |  |
| 197347 | 43789 | Roadkill | Adult | Female | Crawford | Rural | NEG |  |
| 193224 | 43796 | Roadkill | Adult | Male | Scott | Public_land | NEG |  |
| 193355 | 43801 | Roadkill | Adult | Male | Clinton | Rural | NEG |  |
| 193356 | 43801 | Roadkill | Adult | Male | Clinton | Rural | NEG |  |
| 193363 | 43803 | Roadkill | Adult | Female | Scott | Rural | NEG |  |

**SAMPLES INCLUDED**
| Sample ID | Date Collected | Type | Age | Sex | County | Type2 | Rt-PCR Ct value | SARSCoV-2 Line |
| --- | --- | --- | --- | --- | --- | --- | --- | --- |
| 197200 | 43806 | Hunter Killed | Adult | Male | Crawford | Rural | NEG |  |
| 193625 | 43808 | Hunter Killed | Adult | Male | Clinton | Rural | NEG |  |
| 193628 | 43808 | Hunter Killed | Adult | Male | Clinton | Rural | NEG |  |
| 193670 | 43808 | Hunter Killed | Yearling | Female | Clinton | Rural | NEG |  |
| 193671 | 43808 | Hunter Killed | Yearling | Male | Clinton | Rural | NEG |  |
| 193672 | 43808 | Hunter Killed | Adult | Female | Clinton | Rural | NEG |  |
| 193673 | 43808 | Hunter Killed | Yearling | Female | Clinton | Rural | NEG |  |
| 193674 | 43808 | Hunter Killed | Adult | Female | Clinton | Rural | NEG |  |
| 193648 | 43809 | Hunter Killed | Adult | Female | Clinton | Rural | NEG |  |
| 193725 | 43809 | Hunter Killed | Adult | Male | Clinton | Rural | NEG |  |
| 193726 | 43809 | Hunter Killed | Adult | Female | Clinton | Rural | NEG |  |
| 193727 | 43809 | Hunter Killed | Adult | Female | Clinton | Rural | NEG |  |
| 193728 | 43809 | Hunter Killed | Adult | Female | Clinton | Rural | NEG |  |
| 194509 | 43809 | Hunter Killed | Adult | Female | Taylor | Rural | NEG |  |
| 193724 | 43810 | Hunter Killed | Adult | Male | Clinton | Rural | NEG |  |
| 193733 | 43810 | Hunter Killed | Adult | Male | Clinton | Rural | NEG |  |
| 193734 | 43810 | Hunter Killed | Adult | Male | Clinton | Rural | NEG |  |
| 193735 | 43810 | Hunter Killed | Adult | Male | Clinton | Rural | NEG |  |
| 193736 | 43810 | Hunter Killed | Adult | Female | Clinton | Rural | NEG |  |
| 193826 | 43810 | Hunter Killed | Yearling | Female | Clinton | Rural | NEG |  |
| 193828 | 43810 | Hunter Killed | Yearling | Male | Clinton | Rural | NEG |  |
| 193829 | 43810 | Hunter Killed | Adult | Female | Clinton | Rural | NEG |  |
| 197592 | 43810 | Hunter Killed | Adult | Female | Ida | Public_land | NEG |  |
| 194853 | 43811 | Hunter Killed | Adult | Male | Black Hawk | Public_land | NEG |  |
| 198536 | 43811 | Roadkill | Adult | Female | Adair | Rural | NEG |  |
| 194473 | 43812 | Roadkill | Adult | Male | Hardin | Rural | NEG |  |
| 191696 | 43813 | Taxidermy | Yearling | Female | Taylor | Rural | NEG |  |
| 194527 | 43813 | Roadkill | Yearling | Male | Grundy | Rural | NEG |  |
| 198527 | 43813 | Hunter Killed | Adult | Male | Taylor | Rural | NEG |  |
| 197519 | 43814 | Locker | Adult | Female | Cherokee | Public_land | NEG |  |
| 197520 | 43814 | Locker | Yearling | Male | Cherokee | Public_land | NEG |  |
| 197696 | 43814 | Locker | Adult | Male | Cherokee | Public_land | NEG |  |
| 198747 | 43815 | Hunter Killed | Yearling | Female | Crawford | City_bounds | NEG |  |
| 198765 | 43815 | Hunter Killed | Adult | Male | Cherokee | Public_land | NEG |  |
| 198534 | 43816 | Hunter Killed | Adult | Male | Madison | Rural | NEG |  |
| 198549 | 43816 | Hunter Killed | Adult | Female | Adair | Rural | NEG |  |
| 198749 | 43818 | Hunter Killed | Adult | Male | Ida | Rural | NEG |  |

**SAMPLES INCLUDED**
| Sample ID | Date Collected | Type | Age | Sex | County | Type2 | Rt-PCR Ct value | SARSCoV-2 Line |
| --- | --- | --- | --- | --- | --- | --- | --- | --- |
| 198770 | 43825 | Hunter Killed | Adult | Male | Crawford | Rural | NEG |  |
| 190133 | 43829 | Hunter Killed | Adult | Male | Marion | Rural | NEG |  |
| 190138 | 43832 | Hunter Killed | Adult | Male | Warren | Rural | NEG |  |
| 196660 | 43844 | Hunter Killed | Adult | Female | Louisa | Rural | NEG |  |

**Supplementary Table 2.**
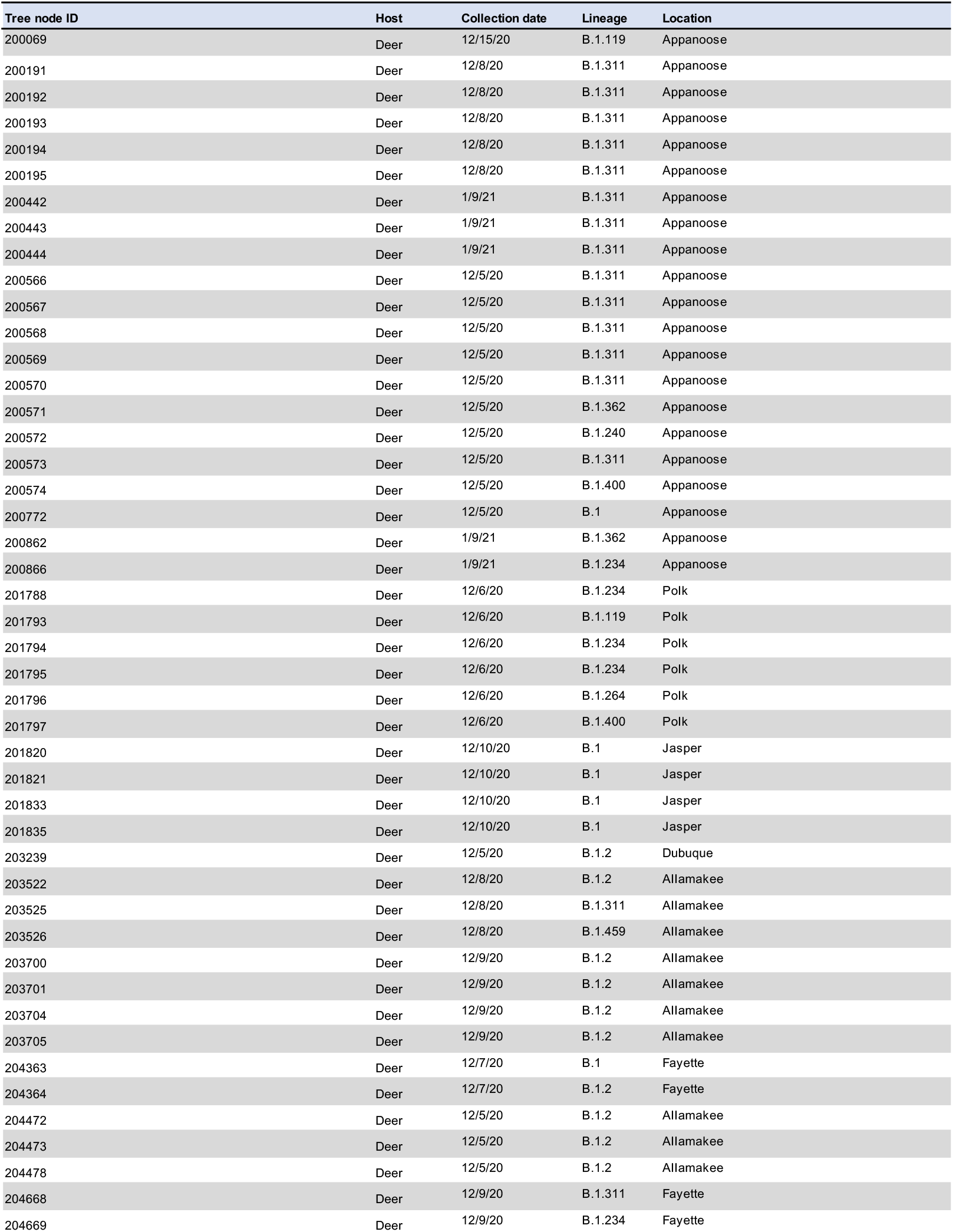
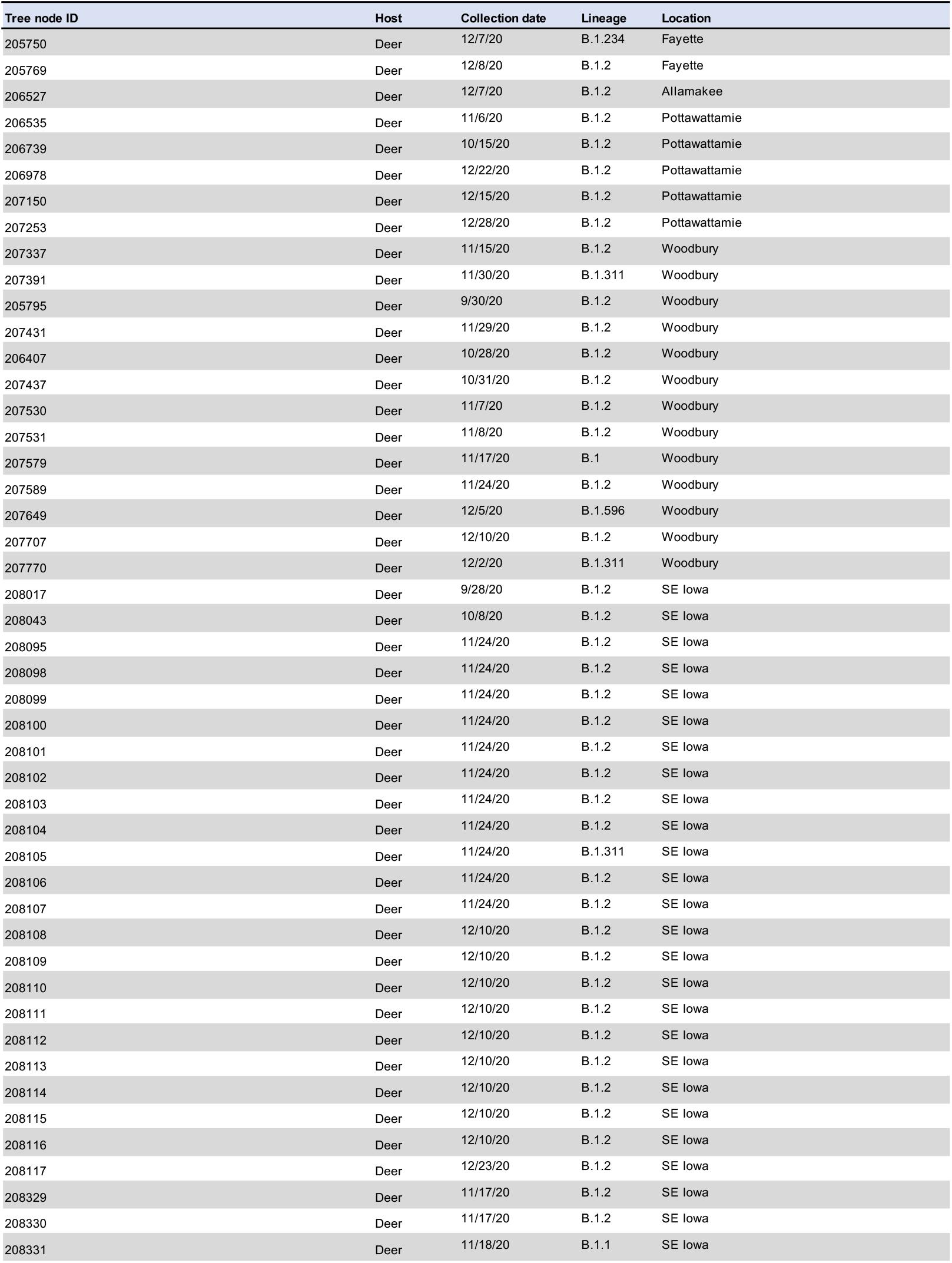
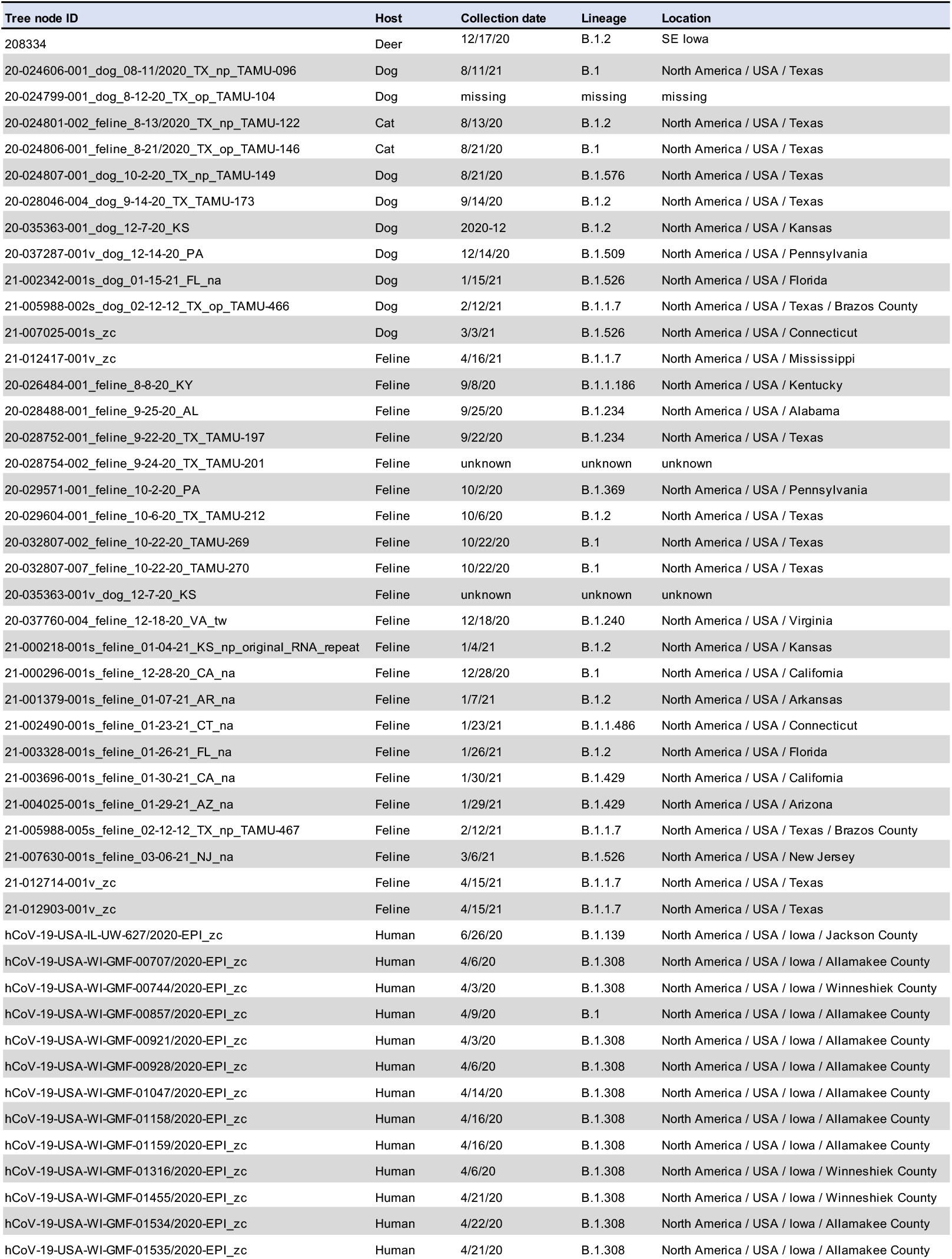
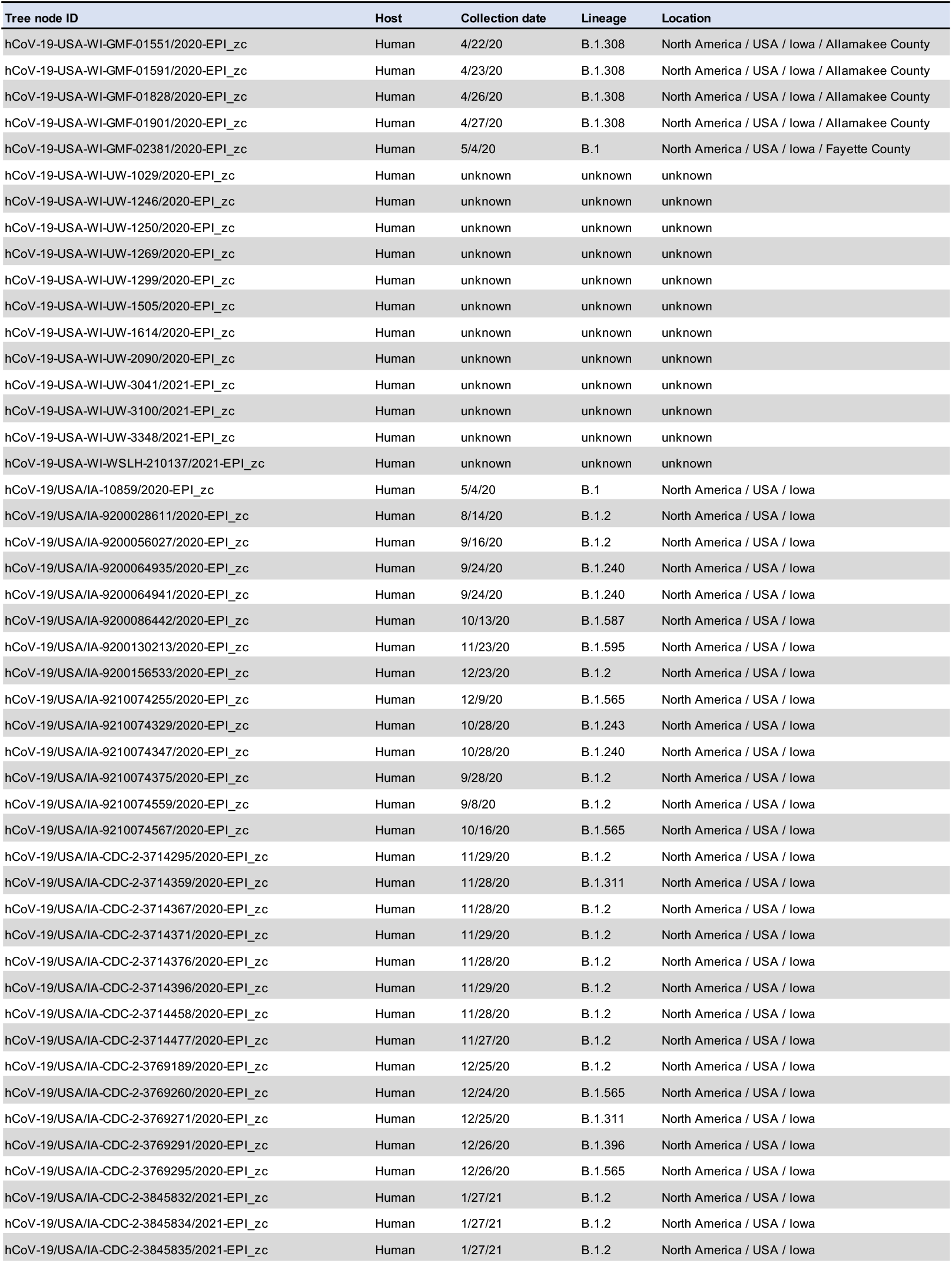
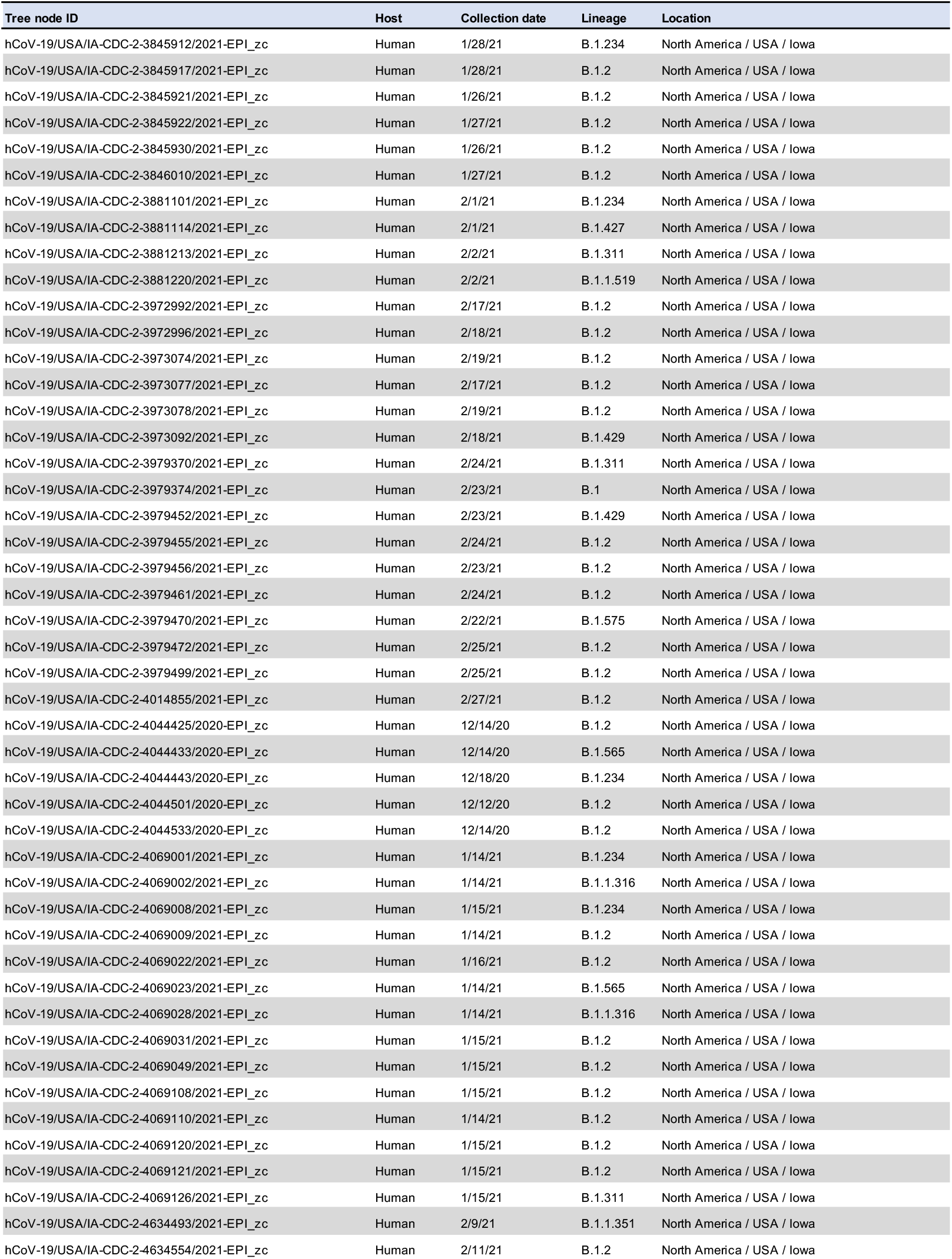
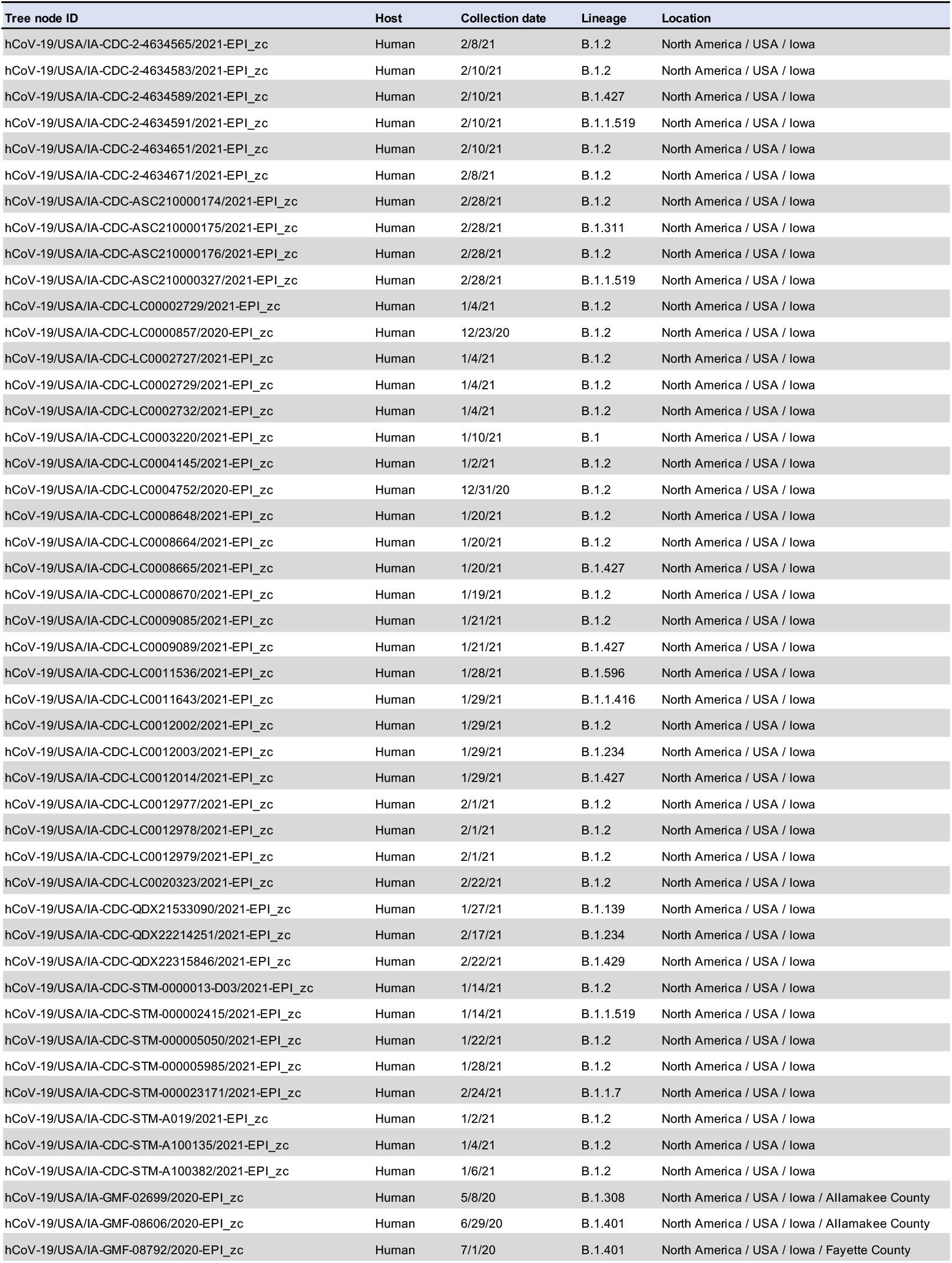
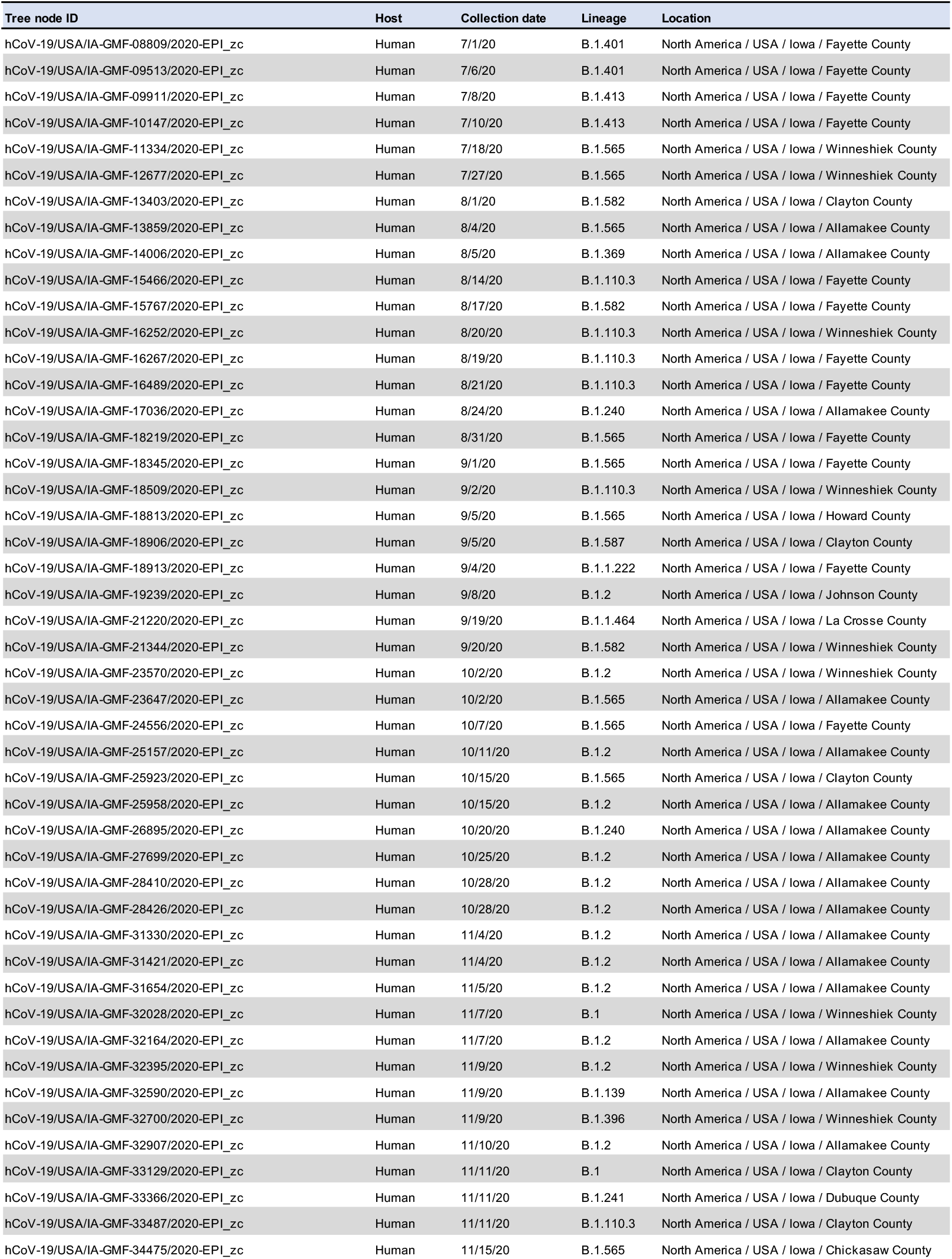
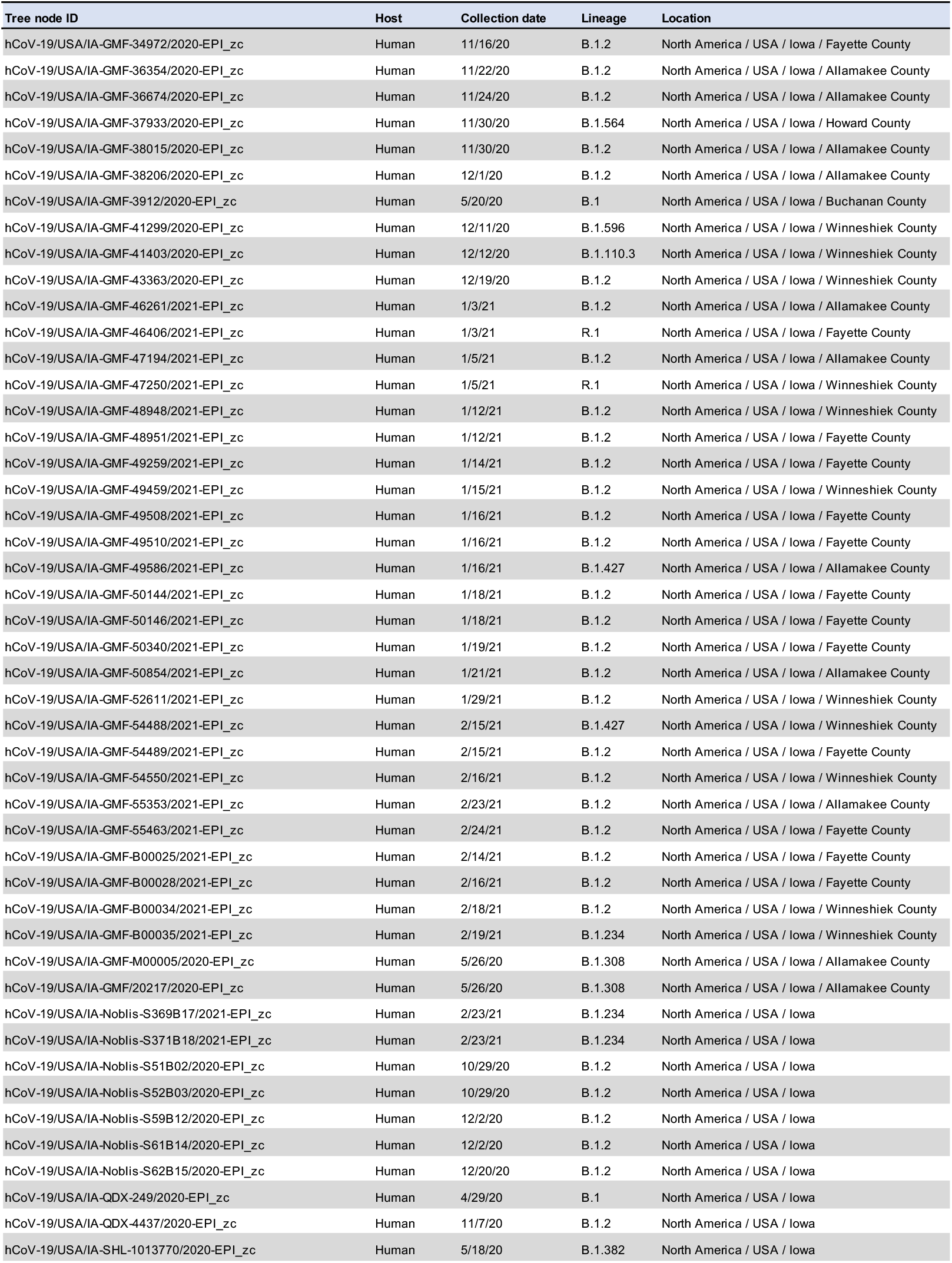
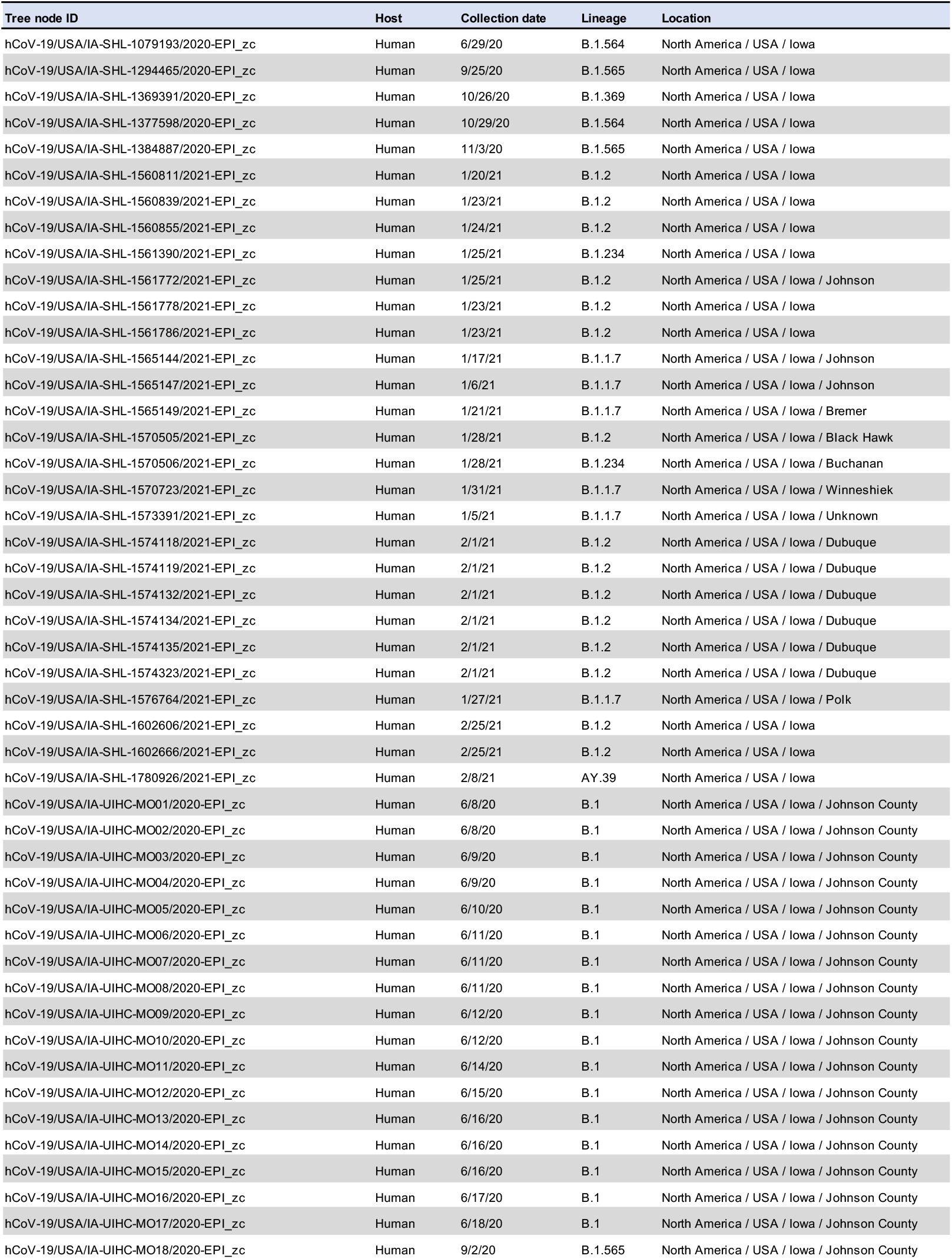
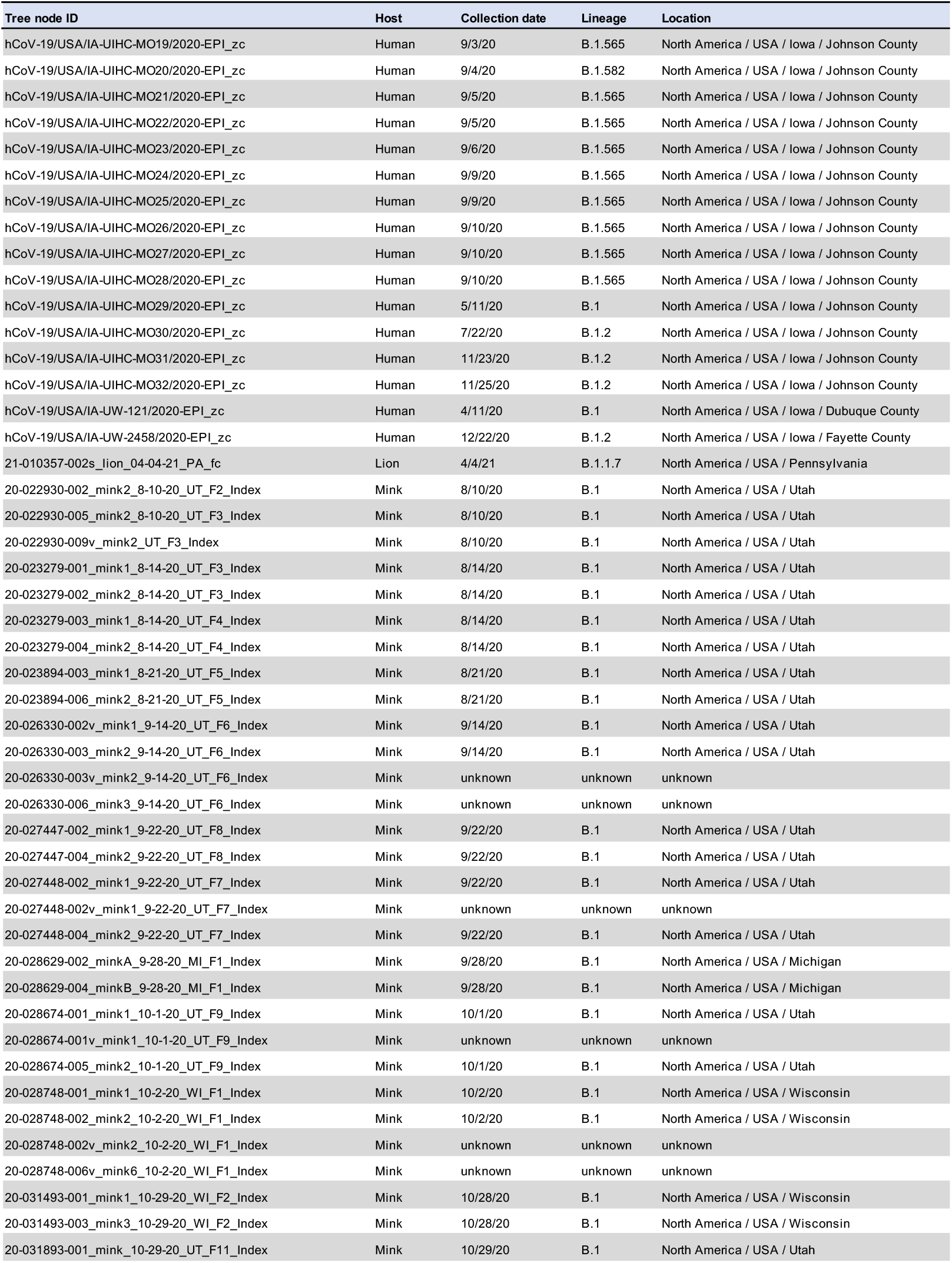
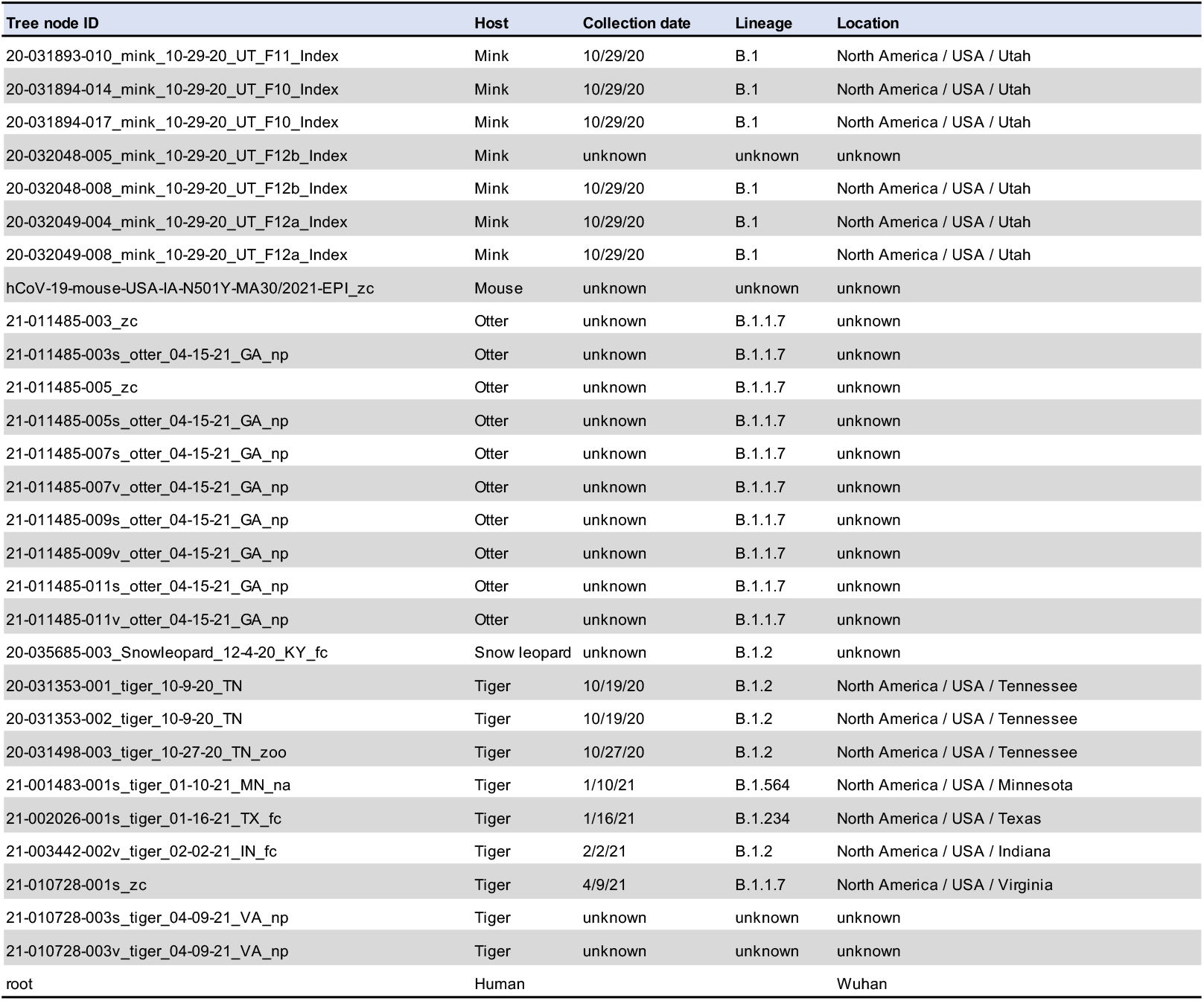

## Notes

### Competing Interest Statement

The authors have declared no competing interest.

